# Regenerated crustacean limbs are precise replicas

**DOI:** 10.1101/2021.12.13.472338

**Authors:** Alba Almazán, Çağrı Çevrim, Jacob M. Musser, Michalis Averof, Mathilde Paris

## Abstract

Animals can regenerate complex organs, yet this frequently results in imprecise replicas of the original structure. In the crustacean *Parhyale*, embryonic and regenerating legs differ in gene expression dynamics but produce apparently similar mature structures. We examine the fidelity of *Parhyale* leg regeneration using complementary approaches to investigate microanatomy, sensory function, cellular composition and cell molecular profiles. We find that regeneration precisely replicates the complex microanatomy and spatial distribution of external sensory organs, and restores their sensory function. Single-nuclei sequencing shows that regenerated and uninjured legs are indistinguishable in terms of cell type composition and transcriptional profiles. This remarkable fidelity highlights the ability of organisms to achieve identical outcomes via distinct processes.

## Main text

The ability to regenerate varies widely among animals. On one extreme of the spectrum, planarians and hydrozoans are thought to be capable of perfect regeneration, using fission and regeneration as means of asexual reproduction (Morgan, 1901). On the other extreme are organisms in which regeneration is lacking (e.g. nematodes) or limited to physiological cell turnover in tissues such as epithelia and blood. Between these two extremes are a wide variety of animals with imperfect regeneration, such as lizards which replace the bony vertebrae of their tails with cartilage (Goss, 1969; McLean and Vickaryous, 2011). In many cases regeneration produces organs of normal appearance that carry subtle defects (Azevedo et al., 2012; Bertozzi et al., 2020; Diogo et al., 2014; Kaars et al., 1984; Lin et al., 2007), but in most instances it is unknown whether cellular composition, detailed morphology and function of regenerated organs are fully restored.

We addressed this question in the crustacean *Parhyale hawaiensis*, an experimental system in which legs typically regenerate within 1-2 weeks after amputation (Alwes et al., 2016; Konstantinides and Averof, 2014). Regenerated *Parhyale* legs are smaller than control legs in the first molt following amputation, but recover in size gradually during subsequent molts (Alwes et al., 2016). On casual inspection, regenerated legs appear identical to uninjured ones in terms of leg segment morphology and the presence of muscles and motor neurons (Konstantinides and Averof, 2014). Despite this, *Parhyale* leg regeneration and leg development differ in their global gene expression dynamics (Sinigaglia et al., 2021).

To survey fidelity in regenerated legs we first focused on the recovery of different types of external sensory organs, exposed as setae on the surface of the legs. These structures provide exquisite markers for assessing the accuracy of regeneration with high almost cellular – precision, because their detailed morphologies and spatial patterns are complex and reproducible among individuals. Like insect sensory bristles, setae are composed of a few cells and take different shapes depending on their type and function (Hartenstein, 2005; Garm and Watling, 2013).

Scanning electron microscopy (SEM) on the three most distal segments of uninjured *Parhyale* legs revealed eight types of setae with distinct morphologies (Figure 1, Suppl. Figure S1). Immunostainings of acetylated alpha-tubulin, which label nerve axons, showed that all the setae are innervated, as expected of sensory organs (Figure 1B and Suppl. Figure S2). Each setal type has a specific distribution on T4 and T5 thoracic legs, either as a single seta in a stereotypic location (e.g. single hooked, curved and plumose setae in the dactylus, single cuspidate seta in the propodus) or in groups of setae arranged in specific spatial patterns (e.g. crowns, combs and ventral arrays made of lamellate and twin setae in the propodus and carpus) (Figure 1A,C, Suppl. Figures S3 and S4). In these latter types, the number of setae varies depending on podomere size (see below, Suppl. Figure S5A). The only exception to these reproducible spatial arrangements is seen with the microsetae, which are spaced on the surface of the cuticle without an apparent conserved pattern (Figure 1A).

**Figure 1.**
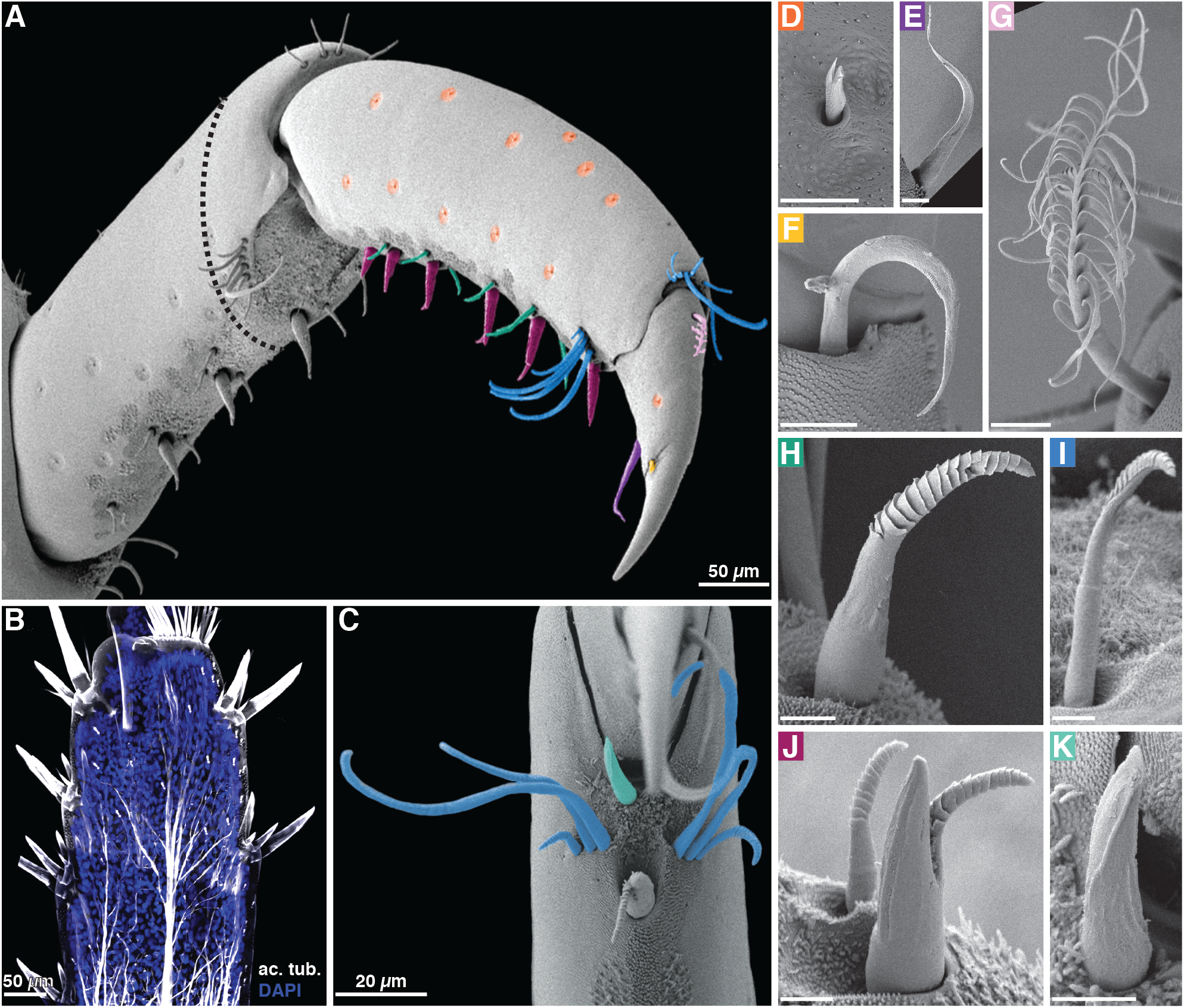
Diversity of external sensory organs in *Parhyale* legs. (**A**) Distal part of an intact *Parhyale* T5 leg, viewed from the posterior by SEM. Different types of external sensory organs (setae) in the propodus and dactylus are highlighted in different colours (details shown in panels D-K). The amputation plane in the setal regeneration experiments is indicated by a black dashed line. (**B**) Acetyl-tubulin staining of the carpus of a T6 leg, showing the innervation of the setae. (**C**) Ventral view of the distal part of the propodus and the dactylus (out of focus) of T5 leg, by SEM, showing the anterior and posterior combs made of type-2 lamellate setae (cyan) and a single cuspidate seta (turquoise). (**D-K**) High magnification view of each type of seta found on the distal part of Parhyale T4 and T5 legs: microseta (D), curved seta (E), hooked seta (F), plumose seta (G), type-1 lamellate seta (H), type-2 lamellate seta (I), twin seta in the foreground (J), and cuspidate seta (K). Panel labels match the colouring of setae in panels A and C. Additional descriptions are given in the Supplementary Text and Figures S1 and S3. Scale bars on D-K, 5 microns.

We then examined whether the diversity, morphology and spatial distribution of these sensory organs are restored during regeneration. To minimise genetic, environmental and physiological effects, we compared the setal patterns between regenerated and control (uninjured contralateral) legs from the same individuals. We found that the detailed morphologies and the spatial distributions of all setal types are unchanged in regenerated legs (Suppl. Figure S1 and Data S1).

Setal types with variable numbers (in the crown, combs and ventral arrays) are less abundant at the first molt post regeneration, but recover during subsequent molts (Figure 2B’, Suppl. Figure S5B). This mirrors leg size, which is initially smaller in regenerated legs but later rebounds to match the size of controls (Figure 2B). Next, we probed the relationship between the number of ventral setae in the propodus and the size of the field in which they develop (see Suppl. Figure S3C). We found that the number of elements increases linearly with podomere length, maintaining a similar spacing of these setae in legs of different sizes (Figure 2A,C). The number of ventral elements found in the regenerated propodus matches that found in similarly-sized uninjured podomeres (Figure 2C).

**Figure 2.**
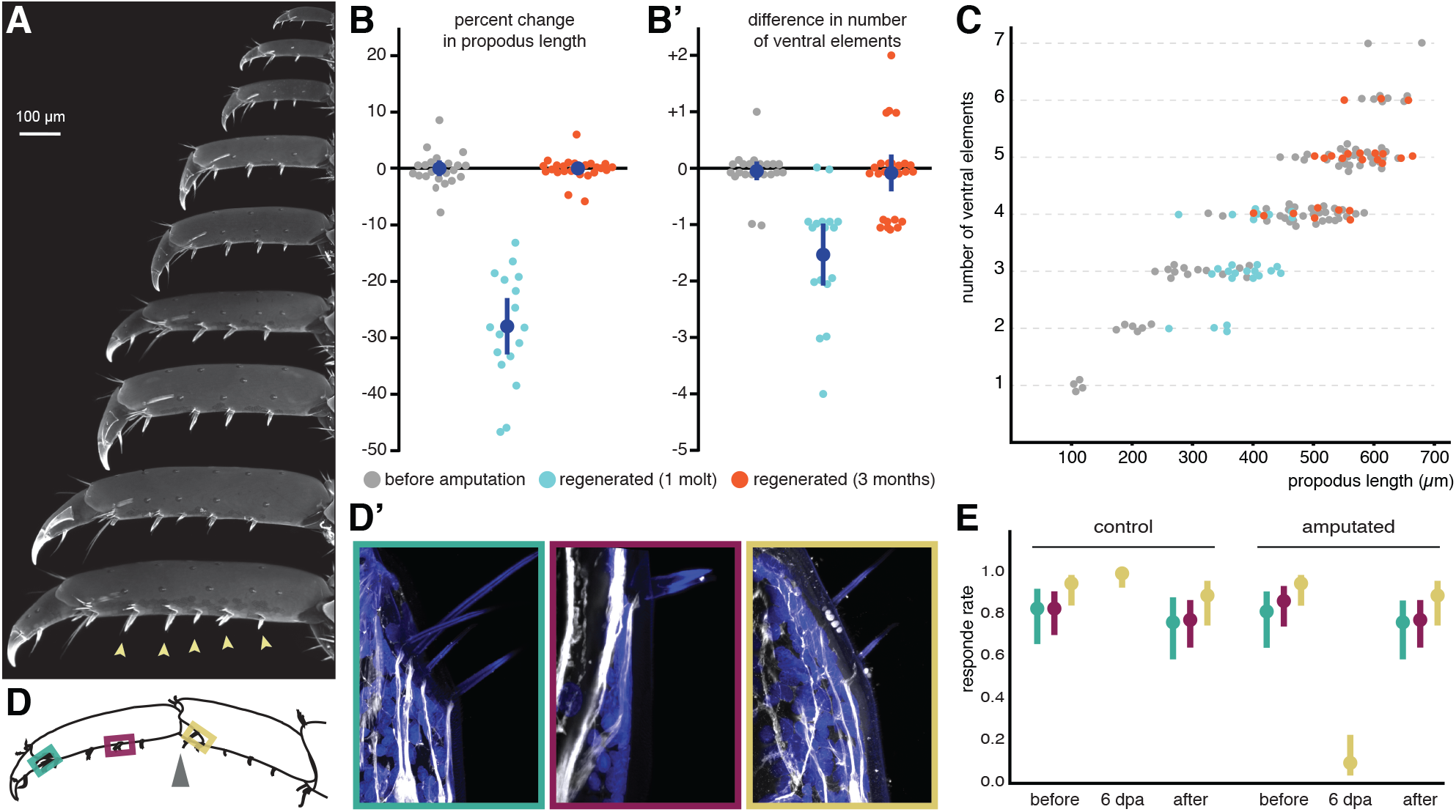
Recovery of sensory organ number, innervation and function. (**A**) The size of *Parhyale* legs varies with age, shown here for the propodus and dactylus of T4 or T5 legs of increasing age (top to bottom). The ventral arrays are made up of a different number of elements (arrowheads) depending on leg size. (**B**) Percent change of propodus length in regenerated versus contralateral uninjured T4 or T5 legs, before amputation (grey), at the first molt post amputation (cyan), and ∼3 months post amputation (red). Newly regenerated legs are smaller than uninjured controls, but their size is fully restored 3 months after amputation. (**B’**) Difference in number of ventral array elements in the propodus of the regenerated and contralateral legs scored in panel B. Newly regenerated legs have fewer ventral arrays, but the number is fully restored within 3 months. Mean values and 95% confidence intervals are shown in dark blue. (**C**) Variation in the number of ventral array elements shown against the length of the propodus in uninjured T4 and T5 legs (grey) and regenerated legs at the first molt post amputation (cyan) and ∼3 months post amputation (red). There is a strong linear correlation between podomere size and the number of ventral arrays. At 3 months post amputation, the number of arrays in regenerated legs is the same as in size-matched uninjured legs. (**D**,**D’**) Acetyl-tubulin staining of comb and ventral array setae after the first molt post amputation, showing that the setae have been re-innervated. (**E**) Mechanosensory responses following mechanical stimulation of comb or ventral array setae in control (uninjured) and regenerated legs of the same individuals (assay shown in Suppl. Movie S1). Legs were amputated at the joint of the propodus and the carpus (arrowhead in panel D). The frequency of response was measured in the same individuals before amputation and after the first molt following regeneration. The comb setae of the propodus (cyan), the ventral setae on the propodus (magenta) and the comb setae of the carpus (yellow) were tested in separate experiments. The combs of the carpus lie just proximal to the amputation plane and are therefore still present at the distal end of the amputated leg stump, in the region of the blastema. When stimulated 6 days post amputation they were found to elicit a very low response rate. Error bars indicate 97.5% confidence intervals.

Immunostainings of acetylated tubulin and a transgenic marker labelling cell outlines showed that the regenerated setae are innervated (Figure 2D,D’, Suppl. Figure S2), suggesting that regenerated setae also recover sensory function. To test this we assayed mechanosensory function by touching comb and ventral array setae (Figure 2D) with a fine probe. This elicited a characteristic escape response (Suppl. Movie S1). Responsiveness was initially impaired in the distal part of the amputated leg stump, but restored to normal levels after the first molt post regeneration (Figure 2E and Suppl. Data S2), indicating that regeneration fully restores mechanosensory function.

Considering the faithful recovery of these microanatomical features, we next investigated whether cellular composition and diversity displayed similar high fidelity in regenerated legs. *Parhyale* legs consist of multiple differentiated cell types, including epidermis, muscles, neurons, tendons, glia, and possibly other, unknown cells. To characterise this diversity we established single-nuclei transcriptional profiling of adult *Parhyale* legs (snRNAseq). This method recovers nuclei from all cell types and overcomes biases in single-cell methods caused by differential dissociation and cell survival, which is especially problematic in animals surrounded by a hard exoskeleton (Grindberg et al., 2013; Lake et al., 2017; Wu et al., 2019).

First, we obtained snRNAseq data for 9,459 nuclei from T4 and T5 legs of adult *Parhyale* (Figure 3A). Reads were mapped to the intronic sequences of annotated genes in order to minimise contamination from ambient cytoplasmic RNA resulting from cell lysis. Dimensionality reduction and clustering analyses separate these nuclei into well-defined clusters, representing cell types or cell states with distinct transcriptional profiles (Figure 3B). We identified clusters corresponding to epidermis, muscle, neurons and blood cells using known cell type marker genes and gene ontology analysis (Figure 3C, Supplementary Text and Suppl. Figure S6A,B). We also tentatively identified clusters corresponding to accessory cells of sensory organs and glia (Figure 3C, Supplementary Text). Unidentified clusters may correspond to known (e.g. tendons) or previously undescribed cell types in crustacean legs (see Suppl. Data S3 for cluster-defining transcripts).

**Figure 3.**
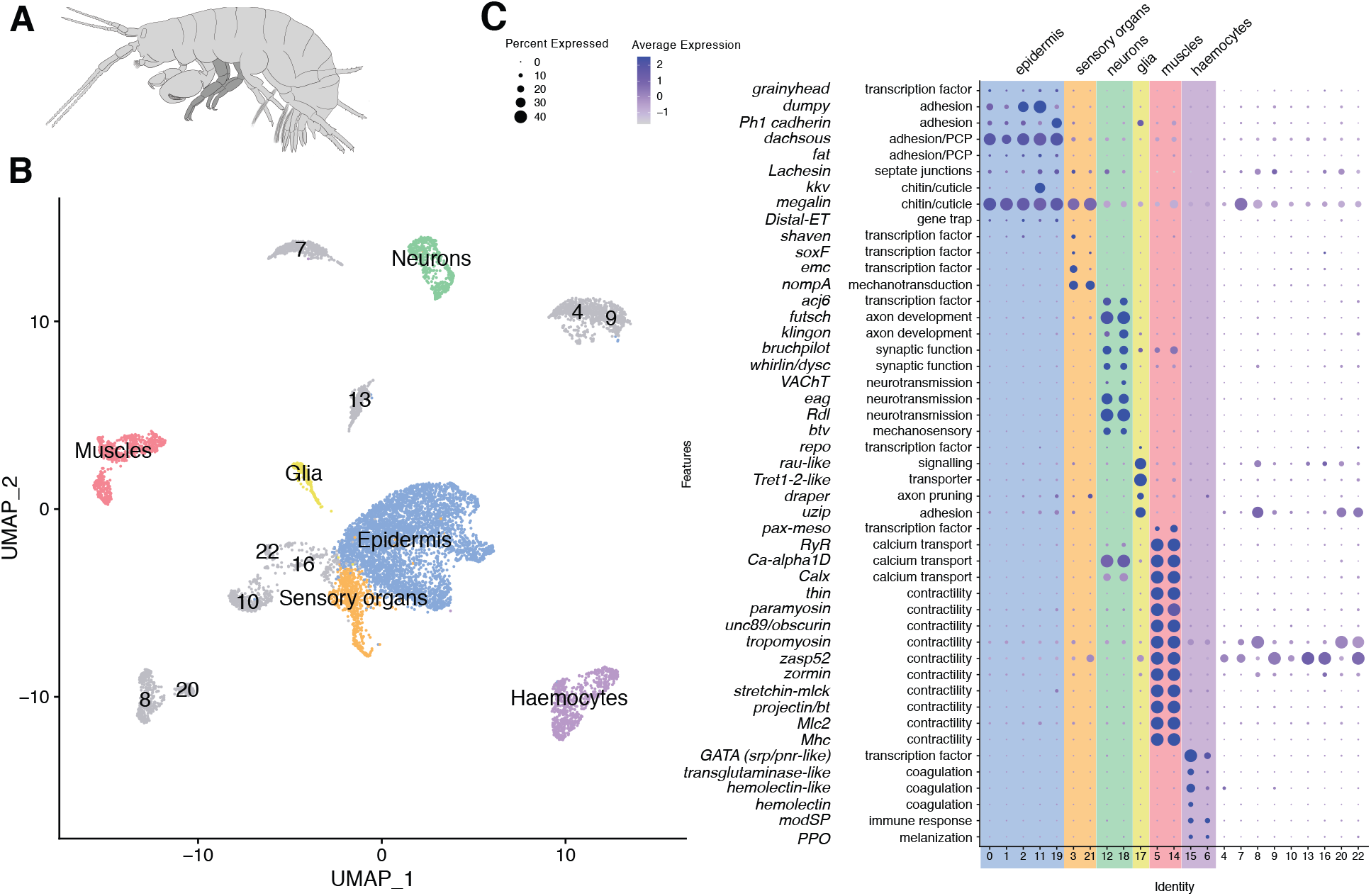
Diversity of cell types in *Parhyale* legs revealed by single-nucleus RNA sequencing. (**A**) Illustration of *Parhyale hawaiensis* adult, with T4 and T5 legs highlighted in dark grey. (**B**) UMAP of snRNAseq experiment on uninjured T4 and T5 adult legs, depicting 9,459 cells clustered based on gene expression. We have identified distinct clusters corresponding to epidermis, muscles, neurons and haemocytes, and putative accessory cells of leg sensory organs and glia (shown in colour, see Supplementary Text); the other clusters are labelled by number. (**C**) Dot plots of selected marker genes for the cell clusters with assigned identities. For each marker we indicate the *Drosophila* or vertebrate orthologue, broad functional category, the average level of expression per cluster (heat map) and the fraction of cells in each cluster expressing the marker (circle size).

To probe cellular composition after regeneration, we performed snRNAseq on regenerated adult legs approximately 2 and 6 months following amputation (see Suppl. Figure S7). To account for genetic, environmental and physiological (e.g. nutritional, circadian, molt- or age-related) effects on the nuclear transcriptomes, we compared snRNA data from regenerated legs (8,925 and 6,668 nuclei from 2 and 6 months post amputation, respectively) with data from contralateral uninjured legs collected simultaneously from the same individuals (8,539 and 7,370 nuclei) (Figure 4A). The analysis was performed *de novo* on the integrated datasets, projecting the data on a new UMAP (Figure 4A-C). In both experiments, the regenerated legs have the same cell clusters as their contralateral controls, including all the clusters corresponding to the major identified cell types, the minor clusters, and those corresponding to yet unidentified cell types (Figure 4B-D). Additionally, we found similar proportions of cells for all clusters in regenerated and contralateral control legs (linear model, ANOVA p-value > 0.08, Figure 4D).

**Figure 4.**
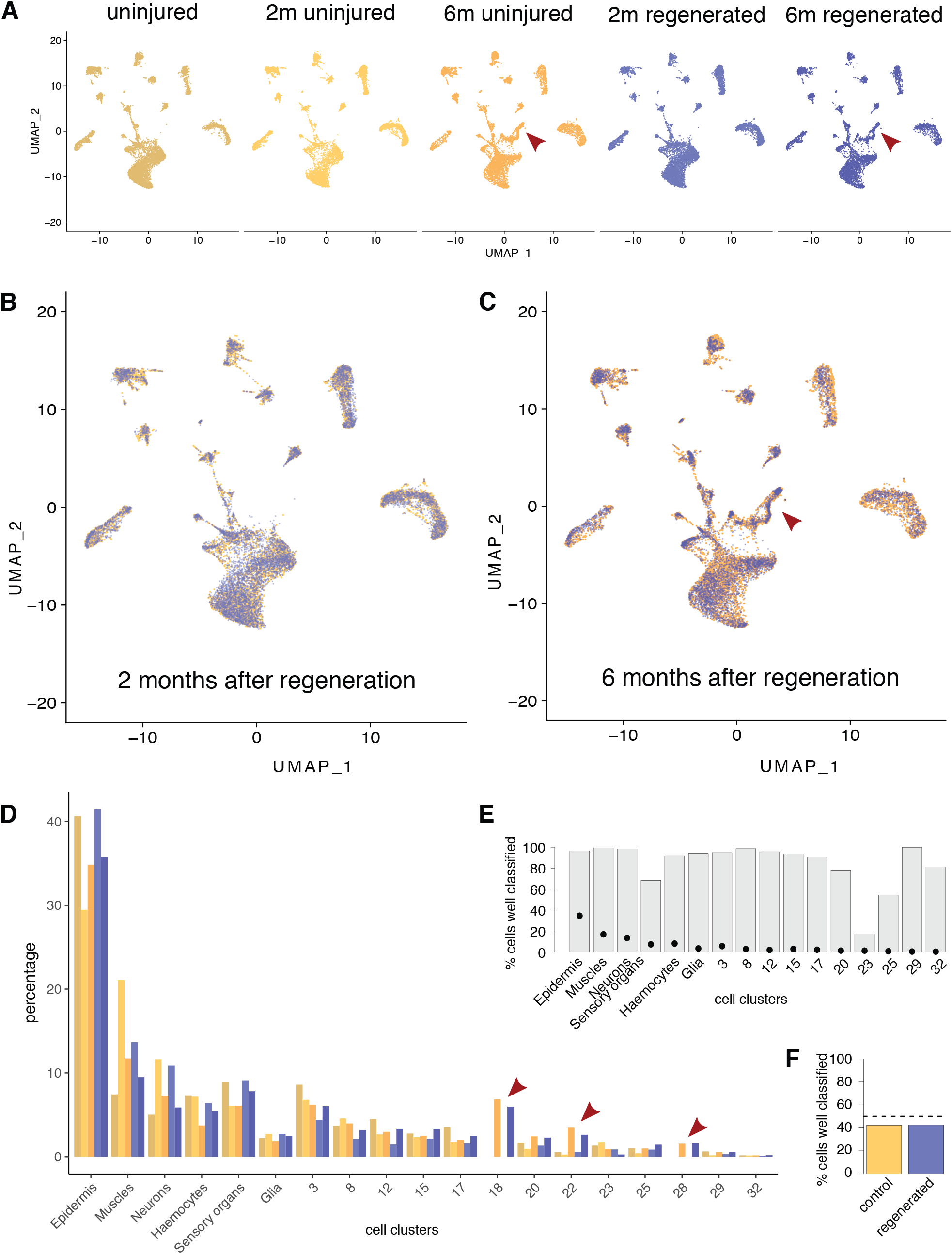
Recovery of diversity and relative numbers of cell types after regeneration. (**A**) Comparison of five snRNAseq datasets (40,961 nuclei), including one dataset from uninjured legs (left, same as in Figure 3B), and datasets from regenerated and control legs sampled ∼2 months and ∼6 months after leg amputation. Samples from uninjured and regenerated legs are highlighted in yellow and blue, respectively. The dimensionality reduction and clustering were performed after integration of the datasets and are independent of those shown in Figure 3B. (**B**,**C**) Overlay of the integrated UMAPs of regenerated and control snRNAseq datasets sampled ∼2 months (B) and ∼6 months after amputation (C). In panel C the epidermal clusters are affected by the expression of molt-associated transcripts in a subset of cells (arrowhead, see Suppl. Figure S11). Within each experiment the same cell clusters are represented in regenerated and control samples. (**D**) Comparison of the relative number of cells recovered per cluster, across all datasets (expressed as % of the total number of cells). Beyond small variations between experiments, we observe no systematic difference between cell proportions in regenerated versus control legs (depicted in shades of blue and yellow, respectively). Cell clusters 18, 22 and 28 (arrowheads in panels A and C and D) are specifically associated with molting (see main text and Suppl. Figure S11). (**E**,**F**) Assessing the effects of cell type (E) and regeneration (F) on the nuclear transcriptome by machine learning (Alquicira-Hernandez et al., 2019) (further detail in Suppl. Figure S13). Predicting whether a cell derives from a control or a regenerated leg based on its nuclear transcriptome is no better than a random guess (dotted line in F). In contrast, the same machine modelling approach can predict a cell’ s cluster classification reliably for most cell types (random guess indicated by dots in E). The lowest prediction accuracy is found for neighboring clusters (for instance 57% of the cells from cluster 23 are predicted as coming from cluster 8; see Suppl. Figures S12A and S13C).

To assess the robustness of our analysis, we tested different methods of data integration (Hao et al., 2021; Welch et al., 2019), including using no integration step, and different read mapping strategies. In the latter case we compared results after mapping RNAseq reads either to introns (which should be found in reads confined to the nucleus) or to the entire genome including unannotated regions, to account for incomplete gene annotation in the *Parhyale* genome (see Materials and Methods). All analyses gave consistent cell clustering and cell abundance measures in regenerated and control legs (Figure 4B-D and Suppl. Figures S8, S9 and S10). In the 6-month experiment, some of these analyses revealed the existence of additional cell clusters in both uninjured and regenerated legs (arrowheads in Figure 4A,D and Suppl. Figures S8 and S10). We found that these correspond to epidermal cell types that express molt-associated transcripts (Suppl. Figure S11, (Sinigaglia et al., 2021)); these cells presumably come from individuals that were close to molting. The detection of this molt-associated cell state confirms that our data integration approach is capable of detecting new clusters.

In addition to cell composition, we also compared the transcriptional profile of each cell cluster between uninjured and regenerated legs (Suppl. Figure S12) and found limited differences that were largely inconsistent across experiments (Suppl. Figure S12B-D). We used machine learning to test whether differences in these transcriptional profiles would be sufficient to distinguish regenerated from control legs. This approach was able to accurately classify different cell types, but could not discriminate between the cells of regenerated and control legs (Figure 4E,F and Suppl. Figure 13).

Overall, our data reveal that the regenerated legs of *Parhyale* are indistinguishable from uninjured legs. This conclusion is based on complementary approaches interrogating the microanatomy, sensory function, cellular composition and molecular profiles of cells. While we cannot exclude the existence of subtle patterning or functional defects in regenerated *Parhyale* legs, at the current level of morphological and molecular resolution these legs appear to be perfect replicas of the original legs prior to amputation.

This remarkable fidelity raises interesting questions about the mechanisms underpinning faithful regeneration in a complex organ. One possible explanation is that the same regulatory networks are deployed to build the organ in embryos and in regenerating adults. Previous studies have supported this notion (Muneoka and Bryant, 1982; Nacu and Tanaka, 2011), however recent work in *Parhyale* and in the sea anemone *Nematostella vectensis* shows that development and regeneration have different temporal profiles of gene expression (Johnston et al., 2021; Sinigaglia et al., 2021). We suggest that development and regeneration could be driven by distinct global regulatory mechanisms, yet incorporate common genetic modules for patterning and cell differentiation that lead to a conserved final outcome.

## Acknowledgements

We thank Francesco Lamanna for advice on snRNA-seq, Marie Sémon for support on statistical analysis, Chiara Sinigaglia for sharing unpublished RNAseq data, Franco Iamunno and the electron microscopy facility of the Stazione Zoologica Anton Dohrn for support on scanning electron microscopy, Benjamin Gillet and Sandrine Hughes for support on NGS sequencing, and the Centre Technologique des Microstructures of the Université Claude Bernard Lyon 1 for help in SEM sample preparation. We are grateful to Evelyn Houliston and Michael Akam for critical comments on the manuscript and to numerous colleagues for additional feedback. This work was supported by the Marie Curie ITN programme EvoCell (H2020-MSCA-ITN-2017 #766053) and the European Research Council grant ReLIVE (ERC-2015-AdG #694918), funded under the European Union Horizon 2020 programme, and by a doctoral fellowship from Boehringer Ingelheim Fonds.

## Supplementary materials

### Materials and methods

#### *Parhyale* culture

Wild type *Parhyale* were raised as described previously (Browne et al., 2005). For all experiments (except Figure S2, see below) we used the wild-type isofemale line Chicago-F (Kao et al., 2016). Several weeks prior to snRNAseq experiments animals were taken from the cultures and kept individually in 6-well plates to ensure uniform conditions and to prevent cannibalism. Similarly, prior to confocal and electron microscopy, animals were kept individually for 3 months in the dark to prevent the build-up of algae. Animals were anaesthetised using 0.02% clove oil.

To examine the innervation of setae (Figure S2) we used transgenic animals carrying the *PhHS>-lyn-tdTomato-2A-H2B-EGFP* transgene, in which lyn-tagged tdTomato (labeling cell outlines) and histone-tagged EGFP (labeling nuclei) are expressed under a heat shock promoter (Alwes et al., 2016).

#### Laser scanning confocal microscopy

For the quantification of setae, legs were fixed for 30 minutes in 3.6% formaldehyde (VWR, #20909.290) in artificial seawater (ASW, specific gravity 1.02), at room temperature. The legs were then washed for 1 hour in ASW, 30 minutes in 50% glycerol in phosphate-buffered saline (PBS), transferred to 70% glycerol and mounted. Cuticle autofluorescence was observed under a Zeiss LSM 800 laser scanning confocal microscope, using the 488 nm excitation laser and a 400-730 nm detection window. The images shown in Figure 2A were cropped manually and juxtaposed on a black background.

For the images shown in Figure S2, the T5 leg of a transgenic animal was amputated on the distal part of the carpus. One day after the molt following amputation, the animal was heat-shocked at 37°C for 45 minutes. One day later the animal was fixed for 15 minutes in 3.6% formaldehyde in ASW and washed thoroughly with seawater. The legs were dissected and mounted in Vectashield mounting medium. The native fluorescence of tdTomato and EGFP was imaged on a Zeiss LSM 800 laser scanning confocal microscope.

#### Scanning electron microscopy

Adult animals of various sizes were selected and the T4 and T5 legs on their right side were amputated at the distal part of the carpus (see dashed line, Figure 1). The uninjured legs on the contralateral side were used as controls. Following the first molt after amputation, the animals were anaesthetised, washed in ASW, fixed for 2 hours in 1% glutaraldehyde (Electron Microscopy Sciences, #16300) in ASW, then washed in ASW and re-fixed for 2 hours in 1% osmium tetroxide (Electron Microscopy Sciences, #19150) in ASW. Animals were then washed in ASW for 1 hour. All steps were carried out at room temperature.

The regenerated T4 and T5 and their contralateral uninjured legs were carefully removed from the fixed individuals, dehydrated by washing in increasing concentrations of ethanol (in ASW) and stored in 90% ethanol. The samples were handled on the proximal parts of the leg in order not to damage the distal structures. Subsequently they were washed 3 times in absolute ethanol (Merck, #1009832500), subjected to critical point drying in a Leica EM CPD300 critical point dryer, mounted on specimen holders, and coated with gold in a Polaron SC7640 sputter coater. Imaging was performed on a JEOL 6700F scanning electron microscope at the electron microscopy facility of the Stazione Zoologica Anton Dohrn, Naples. The images shown in Figure 1A, 1C and S3 were cropped manually and placed on a black background. Colour highlights in Figures 1 and S3 were added as separate layers using the overlay channel option Adobe Photoshop CS6.

#### Testing mechanosensory function

Large adult males were immobilized on standard 60mm Petri dishes from their body using surgical glue (as described in (Alwes et al., 2016)). The setae on T4 and T5 legs were stimulated using a minutien pin (Fine Science Tools, #26002-10) and the animals’ whole-body response was recorded (Suppl. Video S1 and Data S2). Two rounds of experiments were performed under dim light; in the first round, the posterior comb setae of the carpus were stimulated twice (n=22 individuals). After each stimulus the response was recorded and the animals were left to relax for 15 minutes. Following, either one of the T4 (n=10) or T5 (n=12) legs of each animal was amputated. Six days post-amputation, the same setae were stimulated again on both legs, and the animals were left to molt. After molting, they were immobilized again, and the same setae were simulated on both legs. Second round, the animals were glued the same way, and the posterior comb (n=25) and middle ventral array setae (n=28) of the propodus of T4 and T5 legs were stimulated twice with 15 minutes intervals. After either of the legs was amputated and following the molting, the animals were glued, and the same setae were stimulated.

The effect of amputation and subsequent regeneration on the response to mechanical stimulus was assessed using a generalized linear mixed-effects model (R function *glmer* from the package lme4 v1.1-27.1) with a binomial distribution of errors. Sample of origin was tested as a random effect while the leg of origin (T4 or T5), the nature of the leg (amputated or control) and the progress of regeneration (unamputated, 6 dpa, regenerated) were considered as fixed effects. The leg (T4/T5) and the sample of origin had no significant effect on response to stimulus. Amputated legs had a significantly lower response rate 6 days after amputation (p-value < 10-7) whereas the response in fully regenerated legs was not significantly different from the one in unamputated legs. The error bars in Figure 2E indicate 97.5% confidence intervals.

#### Generation of antibodies and immunostaining

A 746 bp fragment from the coding sequence of Parhyale repo (gene MSTRG.29918) was amplified by PCR (primers aaaaatcgataaGATTCAGCACCAGGCAGAG and aaaaatcgatCTGCCGAATTCTTCTTGGAC) and cloned into the *pATH11* plasmid, downstream and in-frame with the *trpE* coding sequence, using the ClaI restriction site. The plasmid was transformed to TOP10 *E*.*coli* competent cells. Transformant colonies were grown in a tryptophan deficient medium, expression of the TrpE-Repo fusion protein was induced by supplementing the medium with 40 μg/ml indoleacrylic acid, and the fusion protein was recovered in inclusion bodies (protocol adapted from (Hoey, 2001)). Mouse immunization and serum collection were performed by Davids Biotechnologie GmbH.

For staining with the Repo antibody (Figure S6C,D), we collected stage S27 embryos (Browne et al., 2005) and T4 and T5 regenerating legs 6 days post amputation. For acetylated tubulin staining in Figure 2D’, the legs were collected 1-2 days after the first molt following amputation. The animals were anesthetized and fixed at room temperature for 10 minutes in 3.6% formaldehyde in ASW. The carpus and propodus of regenerated and uninjured control legs were then dissected and cut in half to improve antibody penetration. The cut legs were re-fixed for 15 minutes in 3.6% formaldehyde in PBS, washed for 1 hour in PBS with 0.1% Triton X-100 (PTx), and incubated for 1 hour at room temperature in PBS with 0.1% Triton X-100, 0.1% sodium deoxycholate, 0.1% bovine serum albumin and 1% normal goat serum (PTxD+). The samples were then incubated for 3 days at 4°C, with mouse monoclonal 6-11B-1 antibody for acetylated alpha-tubulin (diluted 1:1000; Sigma T6793, RRID: AB_477585) or anti-Repo mouse serum (diluted 1:200, see above) in PTxD+. After washing overnight in PTx at 4°C, the samples were incubated with the secondary antibody (anti-mouse IgG Alexa 488, diluted 1:1000; Life Technologies A11001, RRID: AB_2534069) in PTxD+ for 3 days at 4°C. Samples were then washed overnight in PTx at 4°C and mounted in Vectashield mounting medium (Vector Labs, H-1000). The samples were imaged on a Zeiss LSM 800 laser scanning confocal microscope.

#### Nuclear isolation and sequencing

For each snRNAseq dataset we collected a pool of 12-16 thoracic T4 and T5 legs from large adult males. The smallest specimens had a body length of ∼9 mm (estimated age >12 months) and the average had a body length of ∼11 mm (estimated age >18 months). The legs were amputated at the proximal end of the basis (excluding the gills and the coxal plates) using a microsurgical knife (Electron Microscopy Sciences, #72047-30) under anaesthesia in 0.02% clove oil. To compare regenerated and intact legs, T4 and T5 thoracic legs from one side of the animal were amputated and allowed to regenerate and to grow until ∼2 months (8 weeks) after amputation in one pool of individuals, or ∼6 months (24 weeks) after amputation in a separate pool. The contralateral legs were left intact and used as controls. We then collected the sets of regenerated and control T4 and T5 legs by cutting them off at the basis, and dissected them (cut the podomeres lengthwise) in a chilled hypotonic buffer containing 250 mM sucrose, 25 mM KCl, 5 mM MgCl2, 10 mM Tris buffer pH 8.0 and 1 μM DTT and supplemented with 1x protease inhibitor (Sigma, #11873580001), 0.4 U/μl RNasein (Invitrogen, #AM2682), 0.2 U/μl SUPERase-In RNase inhibitor (Invitrogen, #AM2694), 1 mM spermine and 0.1 M spermidine (modified from (Krishnaswami et al., 2016)). The tissue was extruded from the cuticle by gently scraping and peeling using a microsurgical blade and fine forceps (Fine Science Tools, #10316-14 and #11251-20) under a dissecting stereoscope. Triton X-100 was then added to a final concentration of 0.3 % and the tissues were mechanically disrupted by gently pipetting 25 times through a 200 μl filter tip with a cut tip. These steps were carried out in 300 μl of buffer, in the lid of a 5 ml DNA LoBind tube (Eppendorf, #0030108302) kept cold on the surface of a frozen metal plate. The disrupted tissue samples were then transferred to a 1.5 ml DNA LoBind Eppendorf tube (Ependorf, #0030108051), and large pieces of cuticle were let to settle down. The supernatant containing dissociated nuclei was transferred to a new LoBind tube, in which 600 μl fresh hypotonic buffer were added. The nuclei were collected by centrifugation at 500 rcf at 4°C for 6 minutes on a 5424 R Eppendorf microcentrifuge. Centrifugation was kept to a minimum, which resulted in a loose pellet, because longer or faster centrifugations result in lower quality nuclei. The supernatant was discarded and the pellet was washed with 600 μ l 1x Dulbecco’ s phosphate-buffered saline (DPBS) supplemented with 2% bovine serum albumin (BSA). After a second centrifugation, all the supernatant except 25 μL was removed and an equal volume of 3x DPBS with 2% BSA was added to reach a final concentration of 2x DPBS (we observed that nuclei are better preserved but they are more difficult to pellet in 2x DPBS). To remove aggregates, the samples were filtered through 10 μm cell strainer (PluriSelect Mini strainer) to generate single-nucleus suspensions. All the steps were performed in the cold and samples were kept on ice until nucleus capture. The quality of the nuclei was assessed by staining with trypan blue and nuclei counts were made using a Malassez haemocytometer.

Single nucleus RNA sequencing was performed using the 10x Genomics single-cell capturing system. Nuclei suspensions, including 20,000 to 25,000 nuclei, were loaded on the 10x Genomics Chromium Controller following the manufacturer’ s instructions, within 1 hour of preparation. Single nucleus cDNA libraries were prepared using the Chromium Single Cell 3’ Reagent Kit according to the manufacturer’ s instructions (chemistries v3 and v3.1). Libraries were sequenced on an Illumina NextSeq500 (28 bp for R1 and 122 bp for R2) and 248 to 315 million reads were obtained per library. Reads were mapped to the *P. hawaiensis* genome version Phaw_5.0 (see below) using the Cell Ranger Software Suite 6.0.1 (10x Genomics; (Zheng et al., 2017)). Cell Ranger output included lists of features, barcodes and matrices (corresponding to putative genes, cells and read counts, respectively). Subsequent analyses were performed using the R package Seurat v4 (Hao et al., 2021). Quality checks are shown in Figure S14.

#### Genome annotation

The snRNAseq reads were mapped on the *Parhyale hawaiensis* genome assembly Phaw_5.0 (https://www.ncbi.nlm.nih.gov/-assembly/GCA_001587735.2/), in which 19 poorly assembled mitochondrial scaffolds were removed and replaced by the publicly available complete mitochondrial genome (RefSeq NC_039402.1).

To assign reads to genes we used the gene annotation described in Sinigaglia et al. (Sinigaglia et al., 2021). Many genes in this annotation were of undefined or incorrect strandedness. We defined the correct gene strandedness using information from our snRNAseq data (stranded reads from all five snRNAseq experiments) or by identifying putative coding sequences through sequence similarity. First, we compared the total number of snRNAseq reads mapping to opposite strands per gene model. In gene models where at least 80% of reads mapped to one strand (with >20 unambiguously mapped reads in total), we considered that strand as the correct one. In gene models where >20% mapped to opposite strands (or when fewer that 20 reads were recovered) we identified the longest open reading frame on each strand, translated this to a putative protein sequence and used BLAST to search the *Drosophila* and human UNIPROT databases (UP000000803 and UP000005640, respectively) for similar sequences. Strandedness was then assigned based on the best BLAST score (with a minimum threshold of 50). Using this approach we were able to assign strandedness to 40,047 out of 54,699 gene models (the strandedness of 19,334 genes was modified from the original annotation, 13,990 of which using the snRNAseq data and 5,344 of which using coding sequence similarity). For the remaining 14,652 genes, both snRNAseq mapping statistics and BLAST scores were inconclusive, and therefore both strands were included in the gene models (none of those genes had introns). This analysis gave a final list of 69,351 gene models.

As a significant fraction of snRNAseq reads do not map to the exons and introns of the annotated gene models (∼40% of reads), we hypothesized that additional genes could be present in the un-annotated part of the genome. To maximise read mapping and to examine whether non-annotated genes contribute to the snRNAseq signal, we built a secondary annotation that covers the un-annotated part of the genome. For this, we split the intergenic regions of the entire genome into non-overlapping 5 kb fragments (removing fragments below 1 kb, located near genes) and kept the fragments in which we could detect at least 20 unambiguously mapped snRNAseq reads. Strandedness was assigned to each fragment based on the snRNAseq reads mapping to each fragment, as described above (strand assigned if at least 80% of the reads mapped to one strand, both strands included if >20% of the reads mapped to opposite strands). The resulting set included 452,770 annotated intergenic features.

#### Analysis of snRNAseq data

The analysis was performed using the Seurat v4.0.3 scRNA-seq R package, following on the associated documentation (Hao et al., 2021). Genes transcribed in fewer than five cells were removed from the analysis. Cells with fewer than 100 transcribed genes or more than 10% of the genes mapped to the mitochondrial genes, were also excluded from the analysis. Datasets were normalized (Seurat function *LogNormalize*) and variable genes were found using the *vst* method with a maximum of 2000 variable features. Next, the datasets were scaled and principal component analysis (PCA) was performed. A Shared Nearest Neighbor (SNN) graph was computed with 20 dimensions (resolution 1.0) to identify the clusters. Uniform Manifold Approximate and Projection (UMAP) was used to perform clustering and dimensionality reduction. Known markers for epidermis, neurons, sensory organs, glia, muscles and haemocytes were plotted using the Seurat function *DotPlot* (Figure 3C). We identified differentially expressed (DE) genes in each cluster by comparing the expression profiles with those of the rest of the cells using Wilcoxon rank sum-based approach, with a log fold change of >0.25 and a minimum cell percentage of 0.1. The resulting list of DE genes for each cell type was further annotated by identifying putative gene orthologues from *Drosophila melanogaster* and *Homo sapiens* based on reciprocal best BLASTP hits. We functionally annotated our cell clusters by GO term enrichment analysis using the TopGO v2.40.0 package (Alexa A and Rahnenfuhrer J, 2021. topGO: Enrichment Analysis for Gene Ontology. DOI: 10.18129/B9.bioc.topGO) and the database org.Dm.eg.db v3.11.4 (Carlson M, 2019. org.Mm.eg.db: Genome wide annotation for Mouse. DOI: 10.18129/B9.bi-oc.org.Mm.eg.db) on the list of DE genes.

To integrate and compare the five snRNAseq datasets, we used the anchoring approach provided by Seurat. First, we extracted the raw counts from each Seurat object, performed data normalization (function *LogNormalize*) and found variable features independently for each dataset. Then, we performed the identification and integration of the anchors with 20 dimensions, followed by scaling, PCA and the construction of an SNN graph with 20 dimensions (resolution 1). A UMAP was used to perform clustering and dimensionality reduction. Closely related cell clusters were merged and labelled based on the cell-type markers shown in Figure 3C. The relative numbers of cells assigned to each cluster were extracted, calculated from the Seurat object and visualized using the function *barplot* of the ggplot2 v3.3.5 R package. We also performed the same analysis on Seurat with non-integrated data, to test the effect of the anchoring method on the clustering and the recovery of the putative cell types (we obtained similar results, shown in Figure S10).

To examine the impact of the gene annotation in the analysis, we performed the same analysis as described above using the entire genome, including parts of the genome that lack gene annotations (see section on Genome annotation), for read mapping. The two integrated datasets were compared using Clustree v0.4.3 (Zappia and Oshlack, 2018) to visualize the overlap of clusters. The datasets were also integrated using Liger v1.0.0 package (Welch et al., 2019) and SeuratWrappers v0.3.0 to explore the effect of the integration method. The datasets were first pre-processed separately (normalization, finding most variable features, and scaling the data without centering) according to the Liger tutorial. We then calculated the integrative nonnegative matrix factorization (iNMF) with 20 matrix factors to identify the joint clusters and perform quantile normalization by dataset, factor, and cluster, in order to integrate the datasets. Additionally, we ran Louvain community detection (resolution 1) and UMAP on Liger to visualize the clustering of cells graphically. The approach described above was used to merge related clusters.

#### Genes and cell clusters associated with molting

Using an existing dataset (Sinigaglia et al., 2021), we identified 57 genes that are strongly upregulated in *Parhyale* legs 3-4 days prior to molting (DESeq2 (Love et al., 2014), p.adj < 0.001). We used the Seurat function *Dotplot* to compare their expression between cell clusters (Figure S11).

#### Comparing the distribution of cells across cell clusters

We tested whether the proportion of cells recovered in each cell cluster is similar in controls and regenerated legs. We used an ANOVA to test the effect of each of three factors on cell count: which condition each cell is associated with (control/regenerated), which cell cluster it belongs to, and which experiment it was sampled from (0month/2months /6months; all experiments shown in Figure 4A). For the regression, we used a negative binomial distribution, because it captured best the dispersion of our cell count dataset. We used the *Anova* function from the R package *car* with a Fisher test and deviance as error estimate.

We found there was no significant effect of the control/regeneration condition (either as a simple explanatory variable or in interaction with other variables). However the experiment and the cell type of origin had a strong impact on cell count distribution (p-values < 0.001 and < 2.2 × 10-16, respectively).

To further identify differences between experiments and cell clusters, we used the R function *emmeans*. In the 6-month dataset there were significantly more cells in one of the cell clusters associated with molting (cluster 18 in Figure 4D, cluster 12 in Figure S8B). In addition, there were significantly more epidermal cells in the 0-month dataset (compared to the 6-month dataset) and significantly more cells in the muscle and neuronal clusters in the 2-month dataset compared to 0-month or 6-month. However, no statistically significant difference was observed between cells from regenerated legs and controls in any cluster. When mapping reads to the entire genome, epidermal cells are more abundant after regeneration (p-value ∼ 0.01), but this difference disappears if we consider the cluster associated with molting (cluster 12) as part of the epidermal cluster.

#### Differential expression analysis

Differential expression analysis of uninjured versus regenerated legs was performed separately on non-normalized and non-integrated counts of the 2-month and 6-month datasets using the R package Seurat v4.0.3 function *FindMarkers* and the *bimod* method. We chose this method because it gave the most exhaustive list of differentially expressed genes compared to other methods (Wilcoxon test, t test, LR test, MAST), which gave very similar but smaller lists of DE genes. Visualization of gene abundance for the differentially expressed genes (in Figure S12D) was done using the Seurat function *DotPlot*.

#### Prediction of control versus regenerated condition based on transcriptome

In order to test the presence of a transcriptional signature of regeneration, we tried to predict the regeneration state (regenerated versus control) for each cell based on our snRNAseq data using the R package scPred v1.9.2. scPred was designed to predict cluster of origin, but we reasoned that it could be used to predict any given status of a cell. As a control, we also used the same procedure to predict the cluster of origin of each cell (epidermis, muscle, etc.).

The 2-month and 6-month datasets were subsampled, such that the number of uninjured and regenerated cells was identical within each cluster for each experiment: we sampled 7,291 cells for each of the uninjured and regenerated 2-month experiments (a total of 14,582 cells) and 5,694 cells for each of the uninjured and regenerated 6-month experiments (a total of 11,388 cells).

Each dataset was normalized and scaled, and a PCA was computed, using the Seurat functions *NormalizeData, ScaleData* and *RunPCA*.

Training of the machine learning model was done either on the 6-month or the 2-month datasets using the scPred functions *getFeatureSpace* and *trainModel* with the model svmRadial (Figures 4E-F and S13D-E, respectively). We tested other models (e.g. *mda, knn, random forest*) but they gave either similar or worse results on the training set.

Testing was done on the dataset that had not been used for training, measuring the percentage of cells for which a correct prediction was made by the trained model. Baseline values corresponding to the percentage of cells that would be correctly assigned just by chance were calculated as the proportions of cells present in the dataset for each category (for instance 50% and 36% of cells in the 2-month dataset classified as ‘regenerated’ or ‘epidermal’, respectively).

### Supplementary Text

#### Description of different types and groups of setae

We used scanning electron microscopy to survey the surface of *Parhyale* legs, focusing on the three most distal podomeres (dactylus, propodus and carpus) of the T4 and T5 thoracic legs. These two legs show almost identical patterns of setae. On the leg surface we observe the following types of setae, which we categorise following nomenclature introduced in previous studies (Garm and Watling, 2013):

##### Hooked setae (Figures 1F and S1B)

Simple setae with a long thin shaft tapering gradually towards the apex. The shaft has a hooked shape, a series of fine nicks prior to the tapering distal region, and a terminal pore. A single hooked seta is found on the posterior side of the dactylus of T4 and T5 legs.

##### Curved setae (Figures 1E and S1B)

Simple setae with a long twisting shaft, bearing a pore approximately 2/3 along the length of the shaft. A single curved seta is found on the ventral side of the dactylus of T4 and T5 legs.

##### Plumose setae (Figures 1G and S1A)

Setae with a long shaft, bearing two opposed rows of long setules, which give it a feathery appearance. The shaft has no visible pores. A single curved seta is found on the dorsal side of the dactylus of T4 and T5 legs.

##### Cuspidate setae (Figures 1C,K and S1C)

usually consists of a twin seta flanked by type-1 lamellate setae; the element at the distal-most end of Simple setae with a relatively short and stout shaft. The shaft bears longitudinal ridges and has no visible pore. A single cuspidate seta is found ventrally on the distal end of the propodus of T4 and T5 legs.

##### Lamellate type-1 setae (Figures 1H and S1E)

Lamellate setae consist of a smooth shaft bearing a series of lamellae towards the tip. Lamellate type-1 setae have a wider base and a shorter shaft than lamellate type-2, and they bear a terminal pore. Lamellate type-1 setae are organised in ventral arrays (see below) in the carpus and propodus of T4 and T5 legs.

##### Lamellate type-2 setae

(Figures 1I and S1D) are similar to lamellate type-1 setae but tend to have a more slender base, a longer shaft and no pore. Lamellate type-2 setae make up the crown and the anterior and posterior combs (see below) in the carpus and propodus of T4 and T5 legs.

##### Twin setae (Figures 1J and S1E)

Composites of cuspidate and lamellate setae. The cuspidate-like main shaft bifurcates into a branch that resembles the tip of a typical lamellate seta with a terminal pore. Twin setae usually make up the central seta of each element of the ventral array (see below) in the carpus and propodus of T4 and T5 legs. The relative size of the cuspidate and the lamellate components of twin setae can vary (Figure S4); we suggest that these differences reflect a developmental transition from type-1 lamellate to twin setae.

##### Microsetae (Figures 1A,D and S1F)

Very small setae bearing a terminal pore covered by a hood. One side of the shaft has a lamellated appearance. Microsetae are associated with characteristic dimples on the cuticular surface (see figure 1A). They are dispersed on the surface of the leg, with no stereotypic arrangement.

Lamellate and twin setae are clustered in groups on the propodus and carpus of T4 and T5 legs, which were categorised as follows:

##### Crown group

A row of type-2 lamellate setae located dorsally on the distal end of the propodus and the carpus (marked blue in Figure S3). The crown consists of a variable number of setae (see Figure S5 and Data S1).

##### Comb groups

Two clusters of type-2 lamellate setae located ventrally on the anterior and posterior sides on the distal end of the propodus and the carpus, flanking the first element of the ventral array (marked green in Figure S3). The combs consist of a variable number of setae (see Figure S5 and Data S1); anterior combs are sometimes reduced to a single seta.

##### Ventral array

Groups of regularly spaced type-1 lamellate and twin setae, distributed on the ventral side of the propodus and the carpus (marked yellow in Figure S3). Each element of the array the podomere is usually a single type-1 lamellate seta (in juveniles) or a single twin seta (in adults). As suggested earlier, type-1 lamellate setae may mature to become twin setae (Figure S4).

#### Identification of cell types in the snRNAseq data

Cluster analysis on our snRNAseq data identified approximately 23 distinct cell clusters. To identify the cell types represented in these clusters we took three complementary approaches. First, we identified transcripts that are enriched within each cluster and performed a Gene Ontology analysis (Figure S6 A,B). Second, we examined the expression of selected marker genes, whose expression and function are associated with specific cell types throughout the metazoa, or in *Drosophila*, which is the closest extensively studied model organism to *Parhyale* (Table S1). Third, in one case it was possible to use antibodies to observe the localisation of a cluster-specific marker in the legs of *Parhyale* (Figure S6 C,D).

Due to intrinsic limitations, we expected that droplet-based snRNAseq would detect only a small subset of all the mRNAs present in a tissue, and to capture different mRNAs with different efficiency. This is in part because nuclear preparations only capture mRNAs in the transitory phase before their export into the cytoplasm. The residence time of each mRNA in the nucleus varies, e.g. depending on the size of the transcription unit (time to transcribe). A second reason for this differential capture is that standard droplet-based approaches rely on the use of oligo-dT primers to capture the mRNAs and to initiate cDNA synthesis. Nascent transcripts present in the nucleus have not yet acquired polyadenylated tails, so the oligo-dT primers usually work by binding spurious stretches of As along the length of the primary transcript (sometimes within long introns). Thus, the efficiency of detecting a given mRNA depends on its residence time in the nucleus (related to the length of the transcription unit) and the presence of these fortuitous priming sites.

In spite of these limitations, a given transcript should be detected with similar efficiency across all cell types. Therefore, droplet-based snRNAseq allows us to identify cell types/states based on the differential expression of the subset of mRNAs that are efficiently captured.

We focus here on marker transcripts that could be found in our datasets. Orthology assignments are based on reciprocal best BLAST hits with *Drosophila* (for gene names see FlyBase, http://fly-base.org/). The numbering of cell clusters below refers to the analysis (UMAP and dot plots) shown in Figure 3.

##### Epidermal cells

Cell clusters 0, 1, 2, 11 and 19 form a large supercluster of cells (almost 40% of captured cells) expressing overlapping sets of markers. Genes whose functions are associated with the arthropod exoskeleton (chitin synthase *kkv* (Ostrowski et al., 2002)), epithelial adhesion to the exoskeleton (*dumpy*, (Wilkin et al., 2000)), adherens junctions (Ph1 cadherin, (Oda et al., 2005; Sasaki et al., 2017)), planar cell polarity (*fat* and *dachsous*, (Strutt and Strutt, 2021)), septate junctions (*Lachesin*, (Llimargas et al., 2004)) and a transcription factor associated with epidermal repair (*grainyhead*, (Mace et al., 2005)) are specifically expressed in this super-cluster (see Figure 3C). We also find that a gene corresponding to an exon trap expressed in the epidermis of Parhyale limbs (*Distal-ET*, (Kontarakis et al., 2011)) is expressed in this supercluster. Individual clusters may represent different subtypes of epidermal cells (e.g. associated with different parts of the leg) or cell states.

##### Muscles

Gene Ontology analysis provides strong evidence for the identification of clusters 5 and 14 as muscle; GO terms for muscle development and differentiation, mesoderm development, calcium transport and sarcomere assembly are strongly enriched (Figure S6A). Genes with conserved muscle-specific functions, such as genes associated with muscle development (*pox meso*, (Duan et al., 2007)), contractility (e.g. *Mhc, projectin/bt, tropomyosin*, (Schnorrer et al., 2010)) and calcium ion transport (*Calx, Ca-alpha1D, RyR*, (Chorna and Hasan, 2012)), are most strongly expressed in these clusters (see Figure 3C).

##### Neurons

Gene Ontology analysis provides strong evidence for the identification of clusters 12 and 18 as neurons; GO terms for synaptic transmission and membrane potential are strongly enriched (Figure S6B). Genes with neuron-specific functions, including genes associated with neuronal cell type specification (*acj6*, (Clyne et al., 1999)), the formation of axons and dendrites (*futsch*, (Hummel et al., 2000)), synaptic function (*bruchpilot* and *whirlin/dysc*, (Jepson et al., 2014; Kittel et al., 2006)), mechanosensory function (*btv*, (Eberl et al., 2000)), and neurotransmission (*VAChT, eag* and *Rdl* (Dubin et al., 1998; Kitamoto et al., 2000; Stilwell and ffrench-Constant, 1998)), are most strongly expressed in these clusters (see Figure 3C). In *Drosophila* these genes are associated with mechanosensory and/or chemosensory neurons, suggesting that this set is likely to include different types of sensory neurons.

##### Haemocytes

Haemocytes (blood cells) enter *Parhyale* legs through the blood circulation. Cells clusters 6 and 15 express genes that are typically associated with arthropod haemocytes, encoding proteins involved in blood coagulation (related to hemolectin and transglutaminase, (Goto et al., 2003; Lindgren et al., 2008)), melanization (phenoloxidase/PPO, (Sugumaran, 2002)) and immune responses (modSP, (Buchon et al., 2009)) (see Figure 3C). We find that these cells also express specifically a GATA transcription factor and a *mys/itgbn*-like beta integrin, whose orthtologies could not be precisely determined; GATA factors are typically associated with the specification of endodermal tissues and blood (Rehorn et al., 1996) and the beta integrins *mys* and *itgbn* play essential roles in encapsulation, hemocyte migration and phagocytosis in *Drosophila* (Irving et al., 2005; Moreira et al., 2013; Nagaosa et al., 2011). A single-cell RNAseq experiment on whole cells collected directly from blood (A. Almazan, M. Paris and M. Averof, unpublished) has confirmed that these genes indeed are expressed in circulating hemocytes.

##### Putative sensory organ accessory cells

Cells clusters 3 and 21 are associated with the large epidermal supercluster described above. These clusters are enriched for expression of a few genes which are associated with three types of sensory organ accessory cells in *Drosophila* (see Figure 3C): *soxF/sox15* and *shaven/Pax2*, encoding transcription factors that play essential roles in the differentiation of sensory organ socket cells (Miller et al., 2009) and shaft cells (Kavaler et al., 1999) respectively; and *nompA*, which plays an essential role in the sheath cells of mechanosensory organs (Chung et al., 2001).

##### Putative glia

Cell cluster 17 is associated with the expression of the putative orthologue of the insect glial cell marker *repo* (Campbell et al., 1994; Xiong et al., 1994). Using an antibody that we raised against this transcription factor, we find that these cells are closely associated with axons (Figure S6 C,D), as would be expected of glia. The cluster is also enriched in the expression of genes associated with signalling, adhesion, sugar transport and axon pruning in the context of neuron-glia interactions in *Drosophila* (*rau, uzip, Tret1-2, draper*; (Awasaki et al., 2006; Ding et al., 2011; MacDonald et al., 2006; Sieglitz et al., 2013; Volkenhoff et al., 2015)).

**Figure S1.**
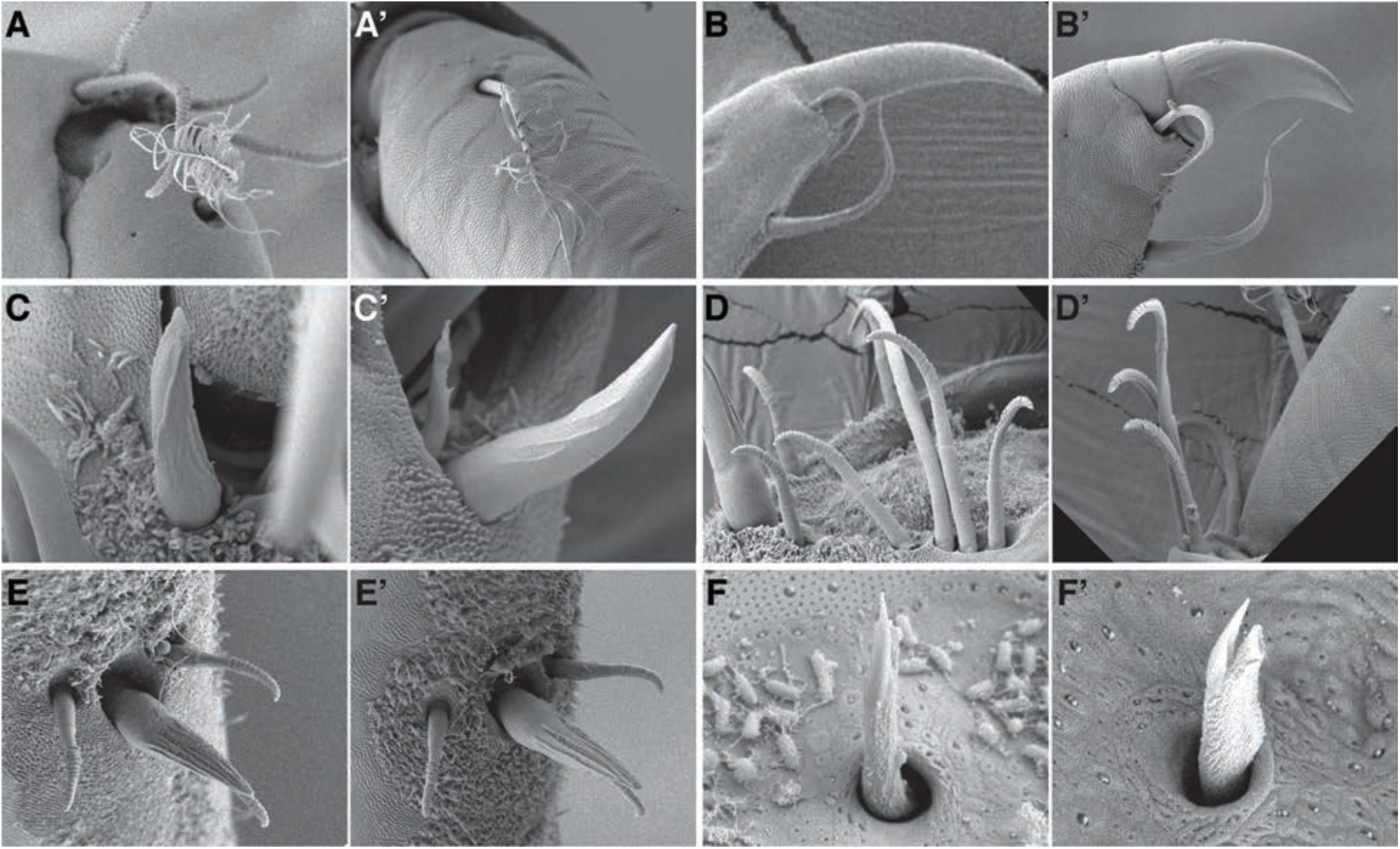
Different types of setae before and after regeneration in T4 or T5 legs. (**A**,**A’**) Plumose setae from uninjured (A) and regenerated (A’) legs. (**B**,**B’**) Distal part of the dactylus bearing hooked and curved setae, from uninjured (B) and regenerated (B’) legs. (**C**,**C’**) Cuspidate setae from uninjured (C) and regenerated (C’) legs. (**D**,**D’**) Type-2 lamellate setae from uninjured (D) and regenerated (D’) legs. (**E**,**E’**) Ventral arrays consisting of type-1 lamellate and twin setae from uninjured (E) and regenerated (E’) legs. (**F**,**F’**) Microsetae from uninjured (F) and regenerated (F’) legs.

**Figure S2.**
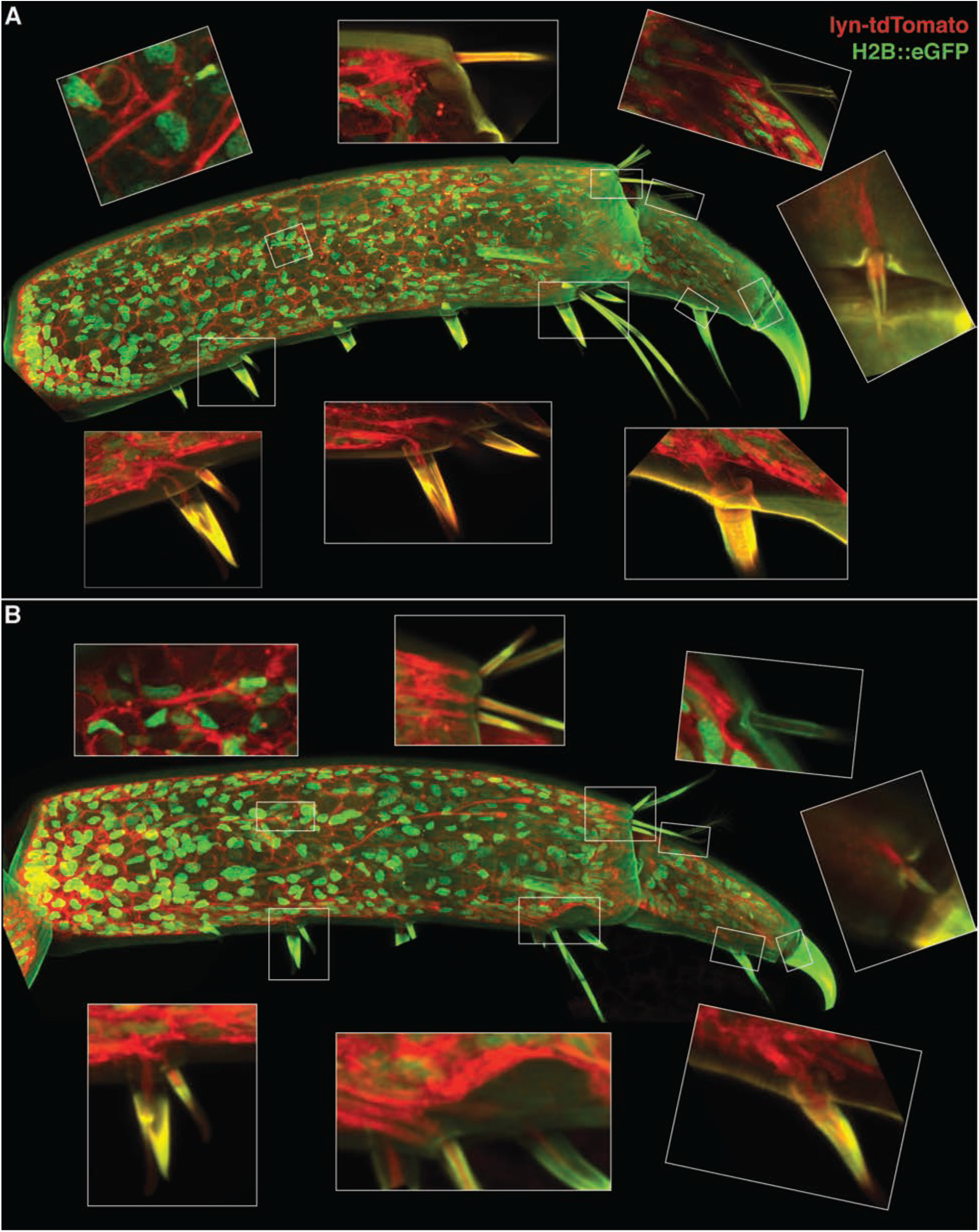
The setae of uninjured and regenerated legs are innervated. Uninjured (**A**) and regenerated (**B**) T5 legs from an individual carrying the *PhHS-lyn>tdTomato-2A-H2B-EGFP* transgene, which expresses lyn-tagged tdTomato (marking cell outlines, in red) and histone-tagged EGFP (marking cell nuclei, in green) in all cells. The legs were imaged 2 days after the molt following regeneration. The tdTomato label allows the processes of the sensory organs to be tracked from the setae on the surface of the epidermis to the main nerve of the leg. In the insets, clockwise from top left: microsetae, type-2 lamellate, plumose, hooked, curved, cuspidate, and lamellate-1 and twin setae. The entire axon tracts of the neurons cannot be seen in this focal plane.

**Figure S3.**
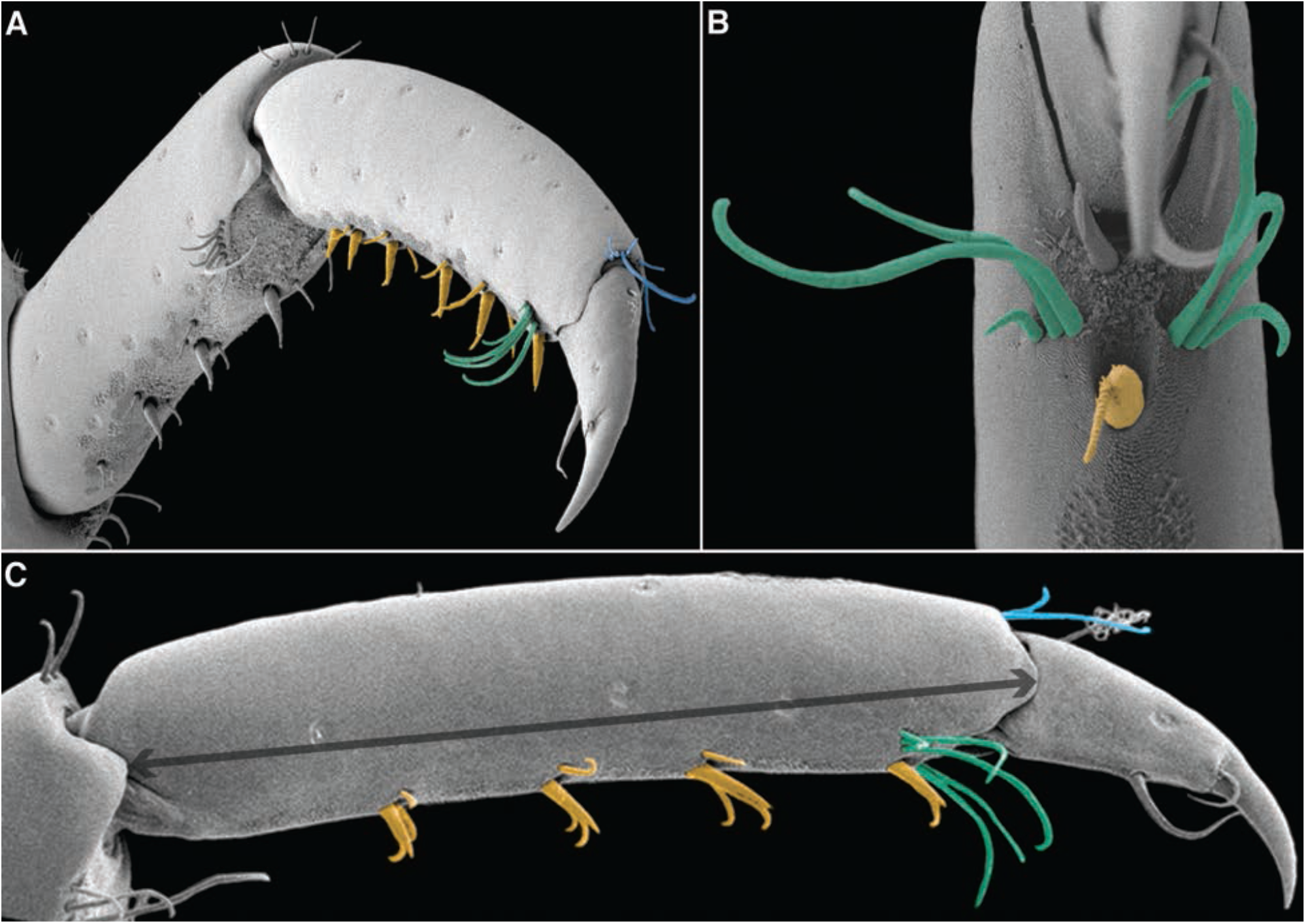
Distribution of the setal groups on the propodus. Scanning electron microscopy images of T5 legs with posterior (**A, C**) and ventral views (**B**). The ventral array (marked with yellow), crown (marked with blue), and comb setae(marked with green) are colored. The length of propodus was assessed by measuring the distance between the distal margins of the propodus and carpus (marked with the arrowed line).

**Figure S4.**
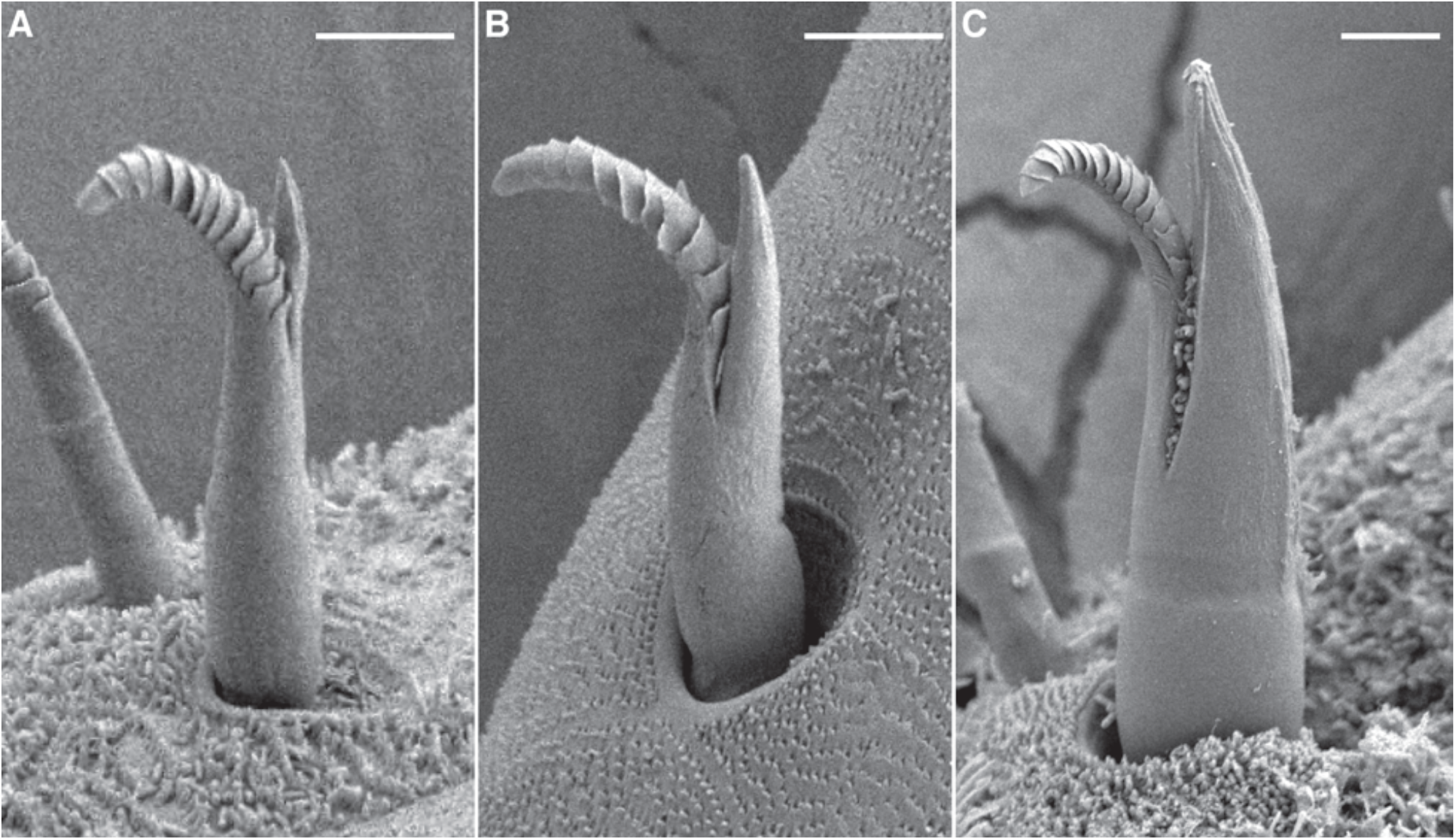
Morphological variation in twin setae. SEM images of twin setae show variation in the relative size of the lamellate and cuspidate components. We propose that these differences represent developmental transitions from type-1 lamellate setae to mature twin setae. The most immature twin setae (**A**) appear very similar to type-1 lamellate setae. More mature types (**B**,**C**) have a more prominent cuspidate branch. The scale bars are 5 microns.

**Figure S5.**
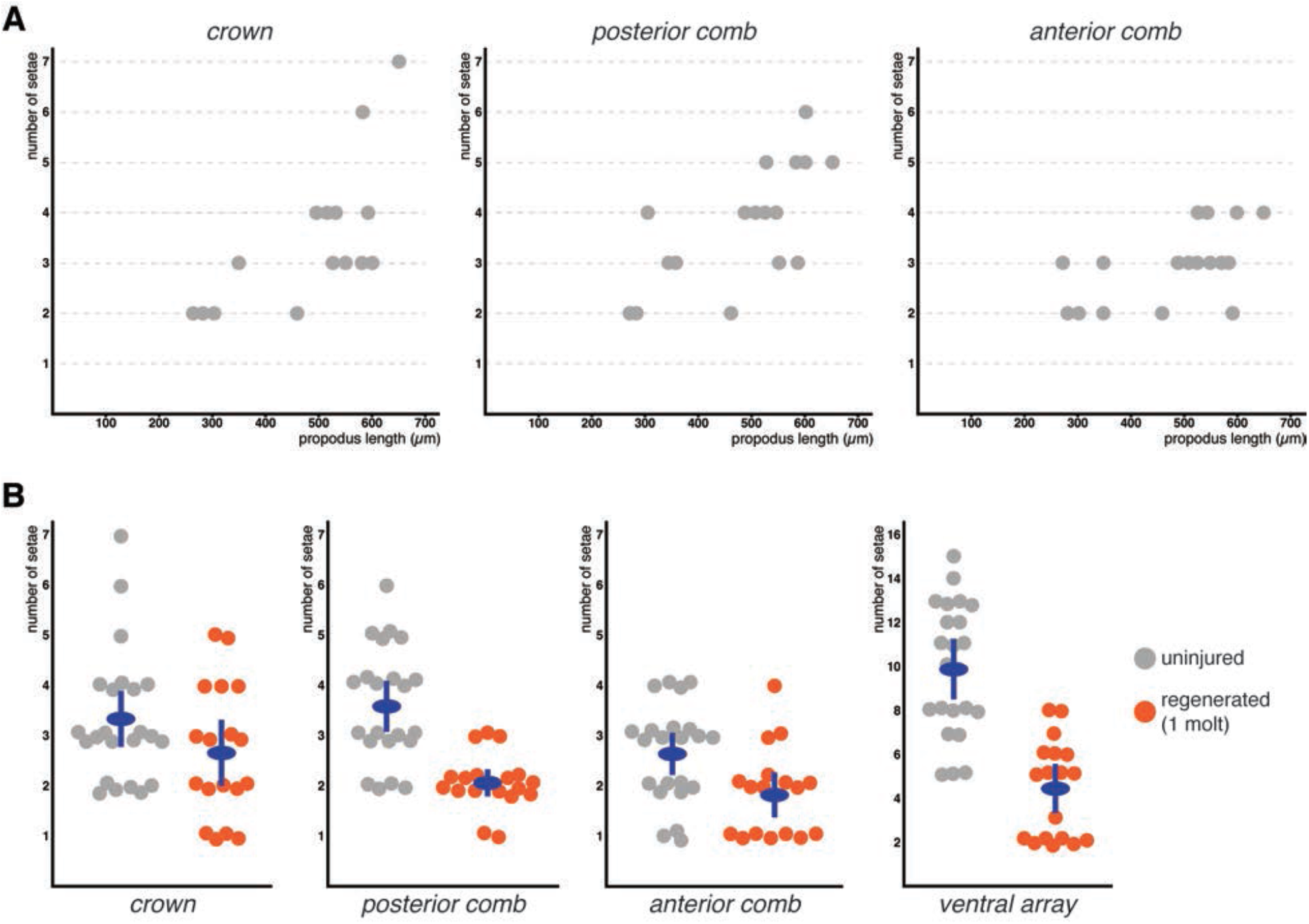
Quantification of setae in the crown, combs and ventral arrays of the propodus. (**A**) Variation in the number of setae in the crown and comb groups shown against the length of the propodus in uninjured T4 and T5 legs. There is a tendency for every group to include more setae as the podomere size gets bigger. (**B**) Quantification of the number of seta in the crown, and comb, and groups, and number of elements in the ventral arrays of the propodus in uninjured and regenerated legs. Each point represents the setae on a single propodus (blue for control, orange for regenerated). Dark blue circles indicate the mean value and the bars mark the 95% confidence interval of the mean. All of the quantifications were performed on legs imaged by scanning electron microscopy (see Data S1 for details).

**Figure S6.**
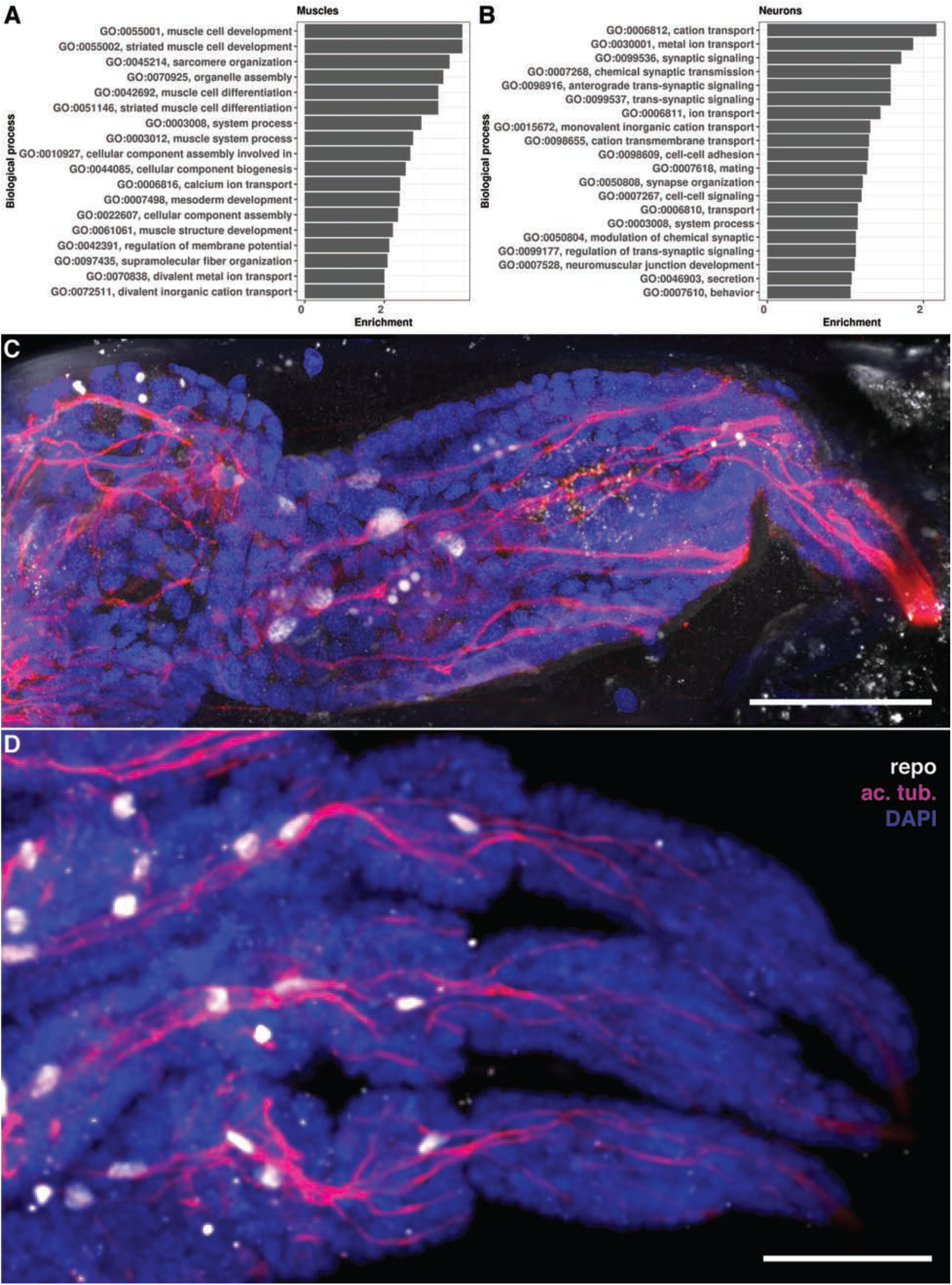
Identification of cell types. (**A**,**B**) Results of GO term enrichment analysis for the putative muscle (A) and neuron (B) cell clusters. (**C**,**D**) Immunostaining of Parhyale embryonic (C) and adult regenerated (D) legs using antibodies raised against the Parhyale Repo protein (white) and acetylated tubulin (magenta); cell nuclei are stained with DAPI (blue). Repo-stained cells are associated with axons. Scale bars are 50 microns.

**Figure S7.**
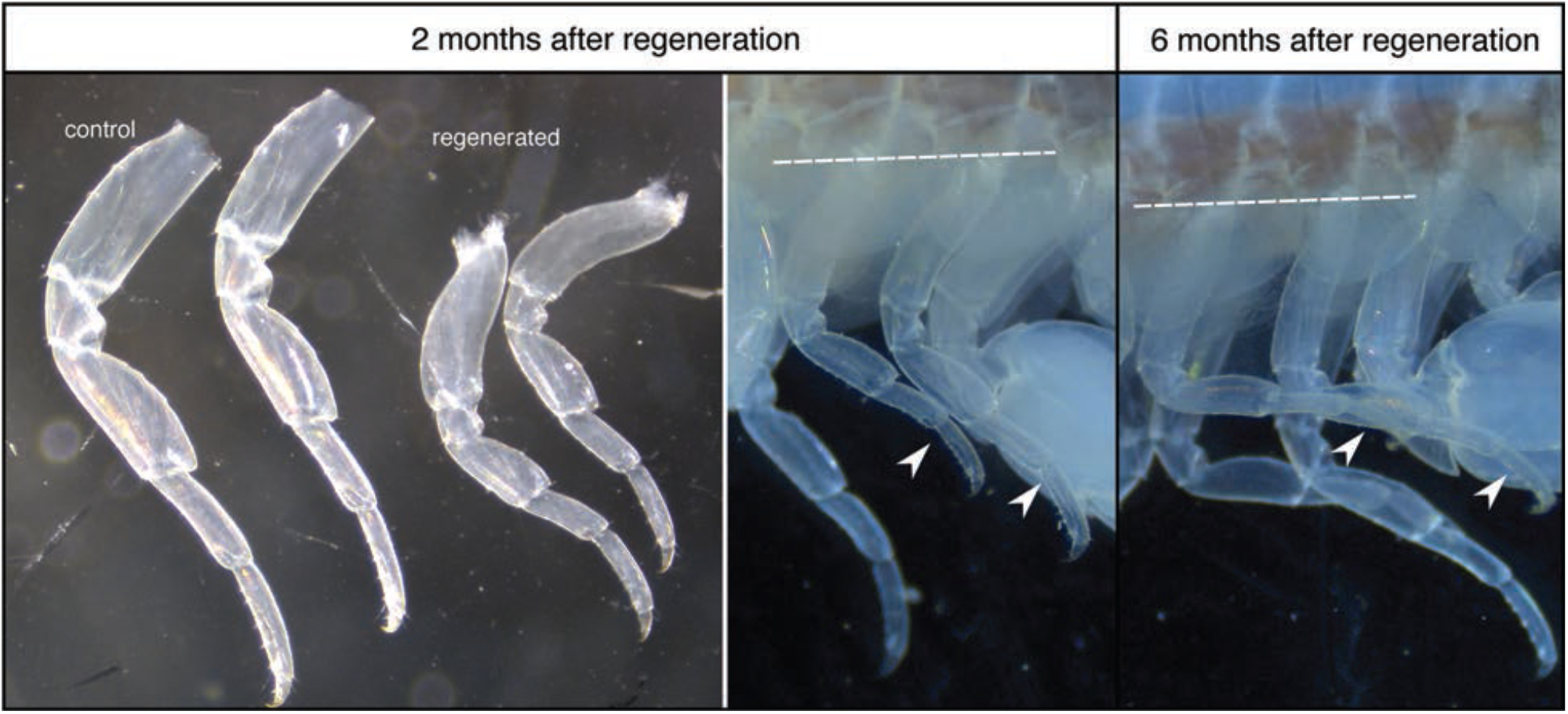
Size of *Parhyale* legs before and after regeneration. The relative size of *Parhyale* T4 and T5 legs is compared between regenerated legs and their contralateral (control) legs approximately 2 and 6 months after amputation. Regenerated legs were smaller than uninjured contralateral legs 2 months after amputation, but had recovered in size by 6 months after amputation (molting interval in these animals was ∼1 month). Regenerated legs are indicated by white arrowheads when imaged on intact animals. Dashed line indicates the amputation plane.

**Figure S8.**
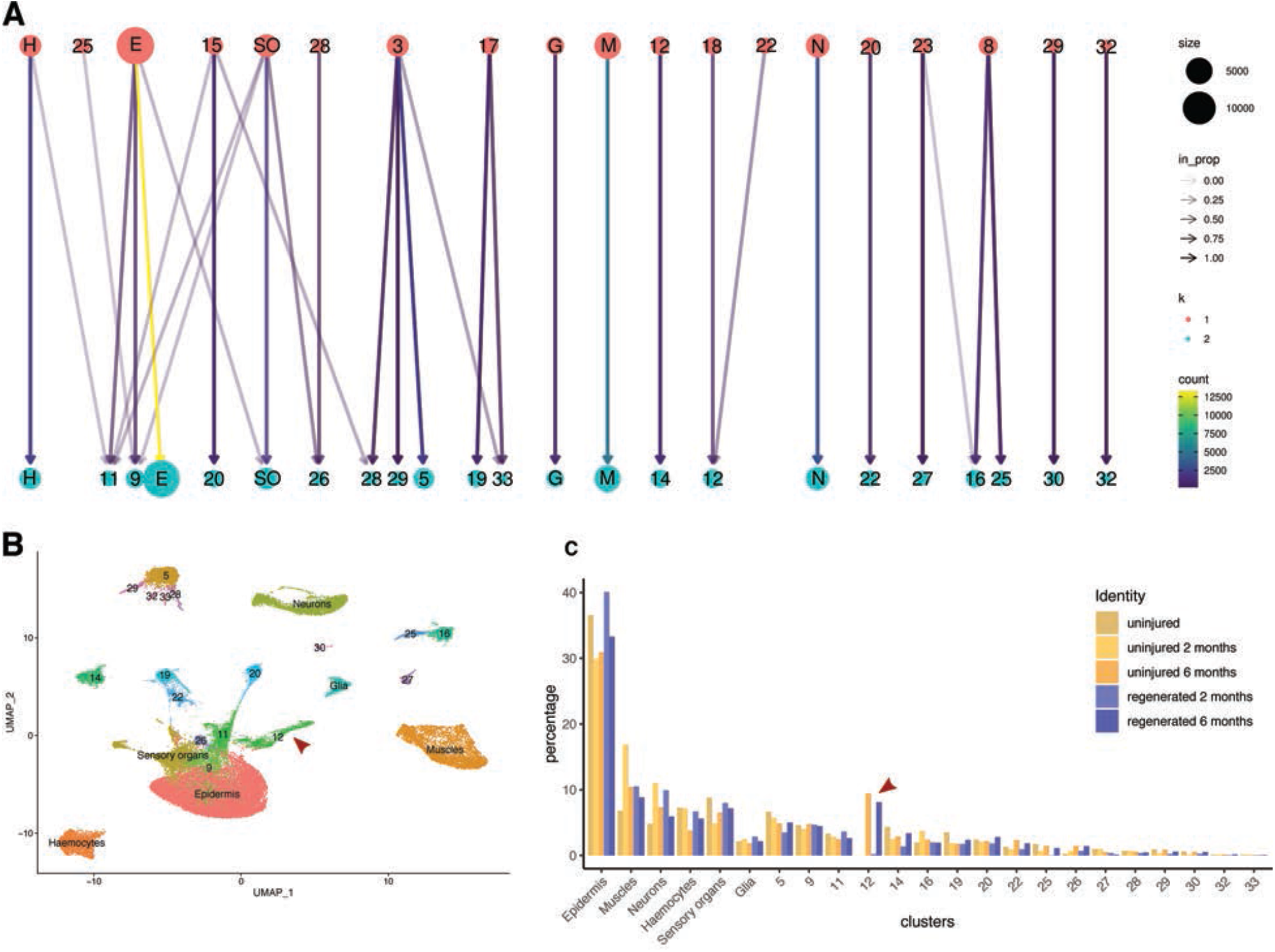
Analysis of the snRNAseq datasets using reads mapped to the entire genome. (**A**) Comparison of the cell clusters identified by mapping the snRNAseq data to introns (top, clusters in red) or to the entire genome (bottom, clusters in cyan). The color of the arrows represents the number of cells shared between clusters, while the intensity represents the fraction of cells that are shared. We observe a good one-to-one correspondence of cell clusters that are well-separated on the UMAP. Cells are reassigned to new clusters particularly among closely-related clusters (e.g. epithelial-related clusters, or clusters 16 and 25 in the whole genome mapping). (**B**) UMAP of the five snRNAseq datasets mapped to the entire genome and integrated in Seurat. The cell clusters corresponding to epidermis, muscles, neurons, haemocytes, and putative clusters for sensory organ accessory cells and glia are labelled as in Figure 3, while the other clusters are labelled by number. (**C**) Comparison of the relative number of cells recovered per cluster, across all datasets (expressed as % of the total number of cells), using reads mapped to the entire genome. As in the previous analysis (Figure 4D), we observe no systematic difference between cell proportions in regenerated versus control legs The arrowheads in panels B and C point to the cluster associated with molting.

**Figure S9.**
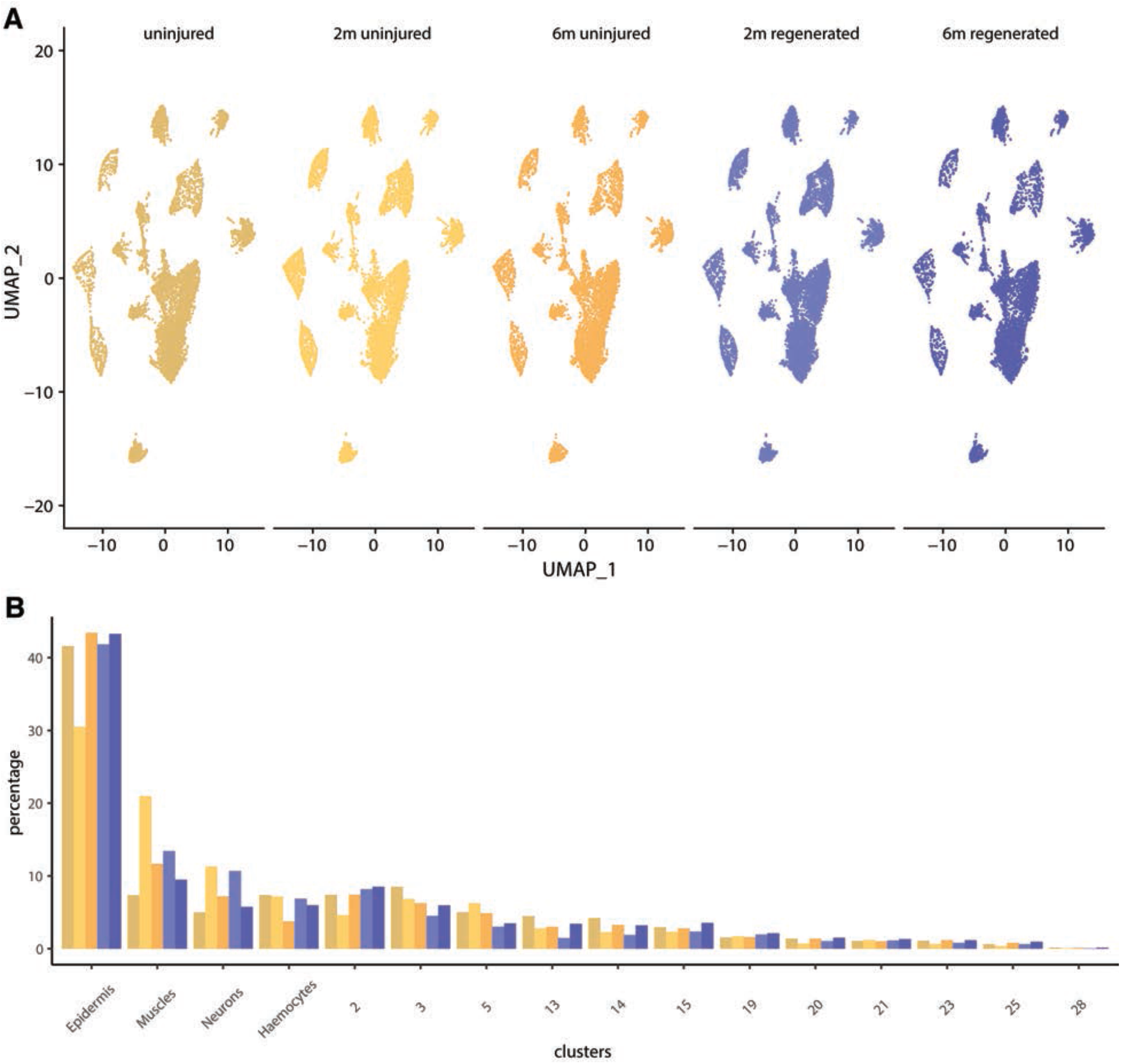
Data integration using integrative non-negative matrix factorization (iNMF, Liger package). (**A**) The five snRNAseq datasets (40,961 nuclei) were integrated using the Liger package and plotted on the same UMAP coordinates. They are displayed using the same colour code as in Figure 4A.(**B**) Comparison of the relative number of cells recovered per cluster, across all datasets (expressed as % of the total number of cells), using iNMF in Liger. As in the previous analyses (Figure 4D and S8C), we observe no systematic difference between cell proportions in regenerated versus control legs.

**Figure S10.**
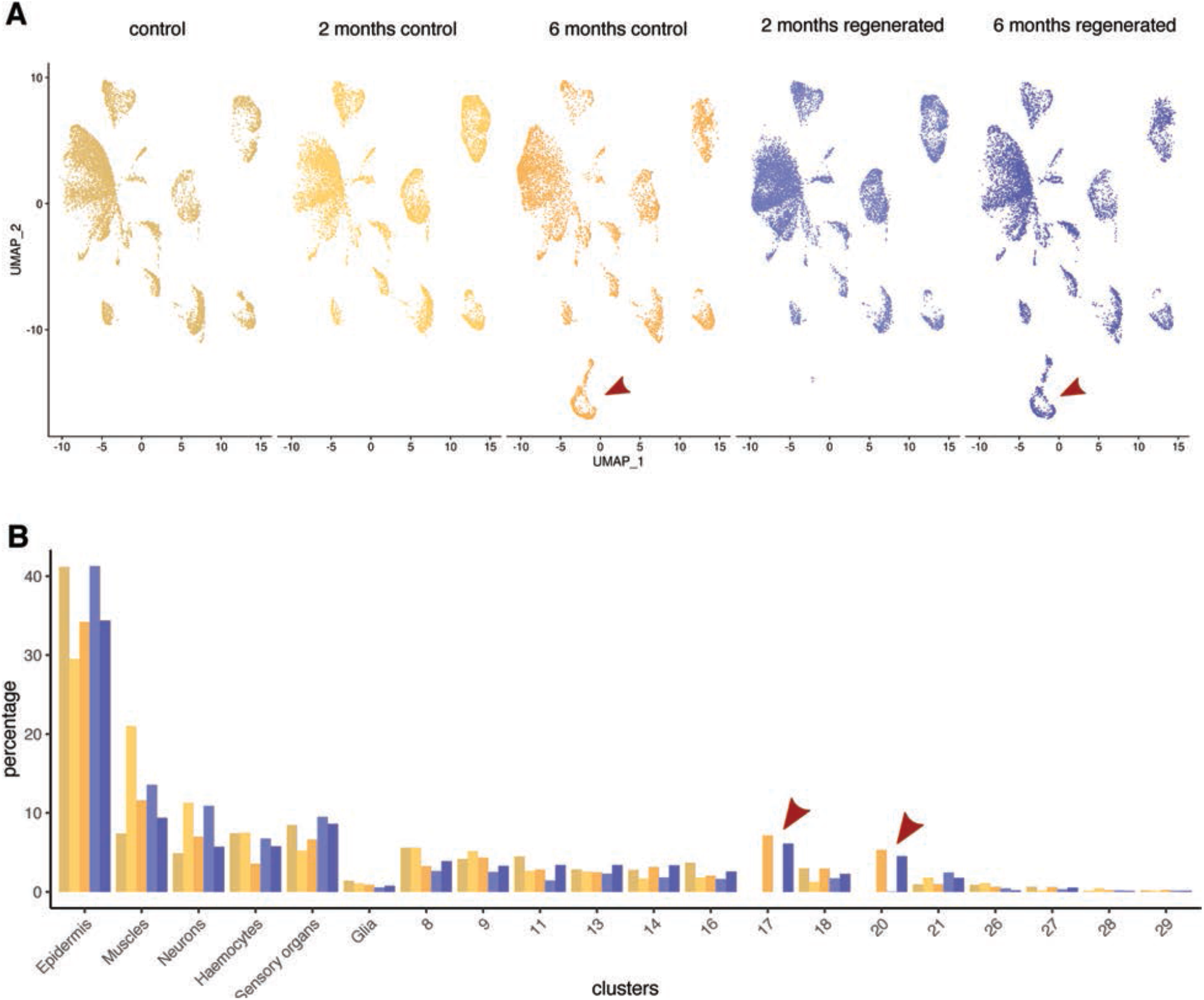
Data aggregation using Seurat. (**A**) The five snRNAseq datasets (40,961 nuclei) were aggregated using the Seurat package and plotted on the same UMAP coordinates. They are displayed using the same colour code as in Figure 4A. (**B**) Comparison of the relative number of cells recovered per cluster, across all datasets (expressed as % of the total number of cells), using data aggregation in Seurat. As in the previous analyses (Figure 4D, S8C and S9B), we observe no systematic difference between cell proportions in regenerated versus control legs. The red arrows indicate the cell clusters associated with molting in the 6-months dataset.

**Figure S11.**
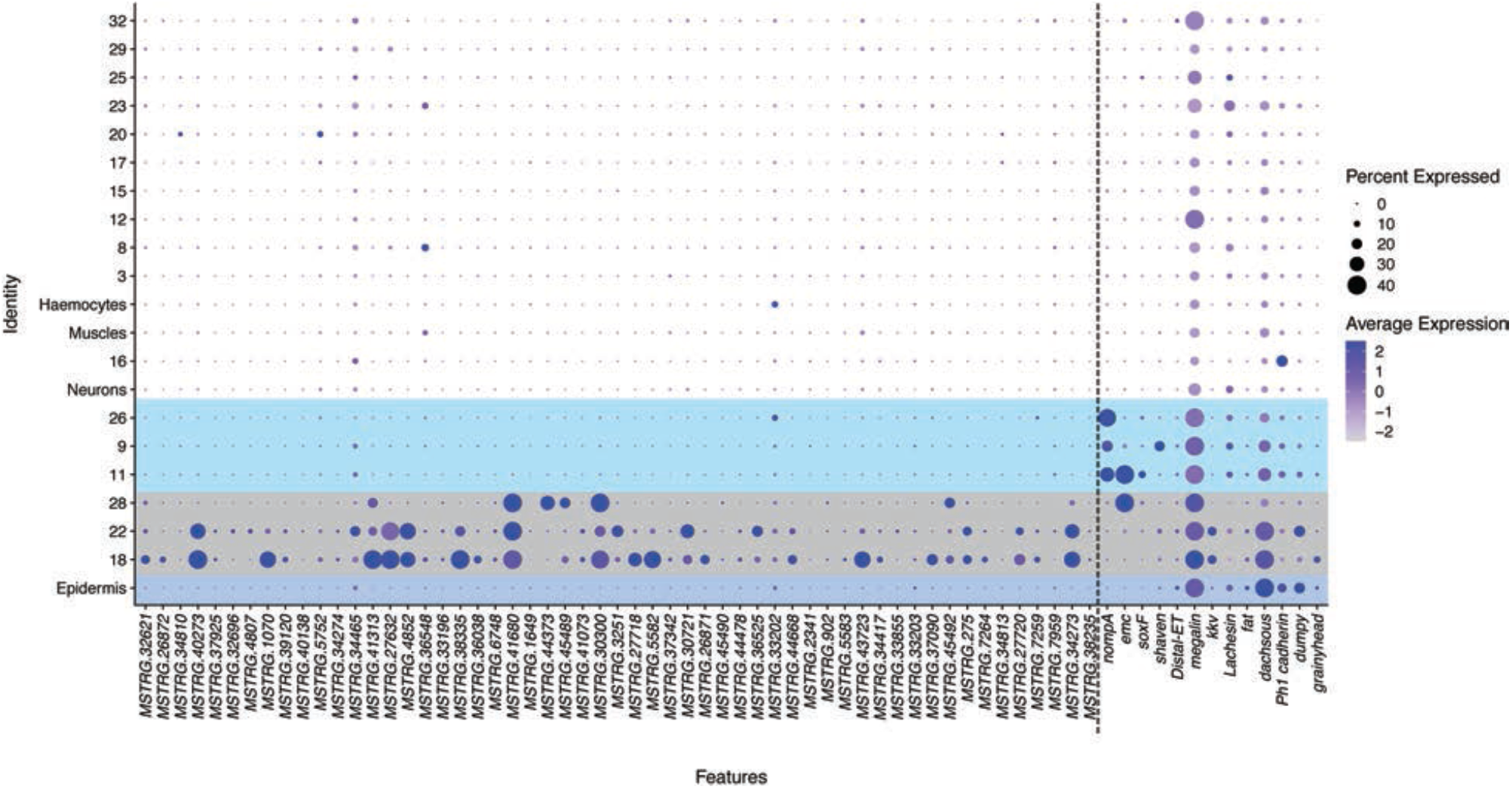
Clusters 18, 22 and 28 express genes that are specifically upregulated during molting. Dot plots of marker genes that are specifically upregulated 3-4 days prior to molting (Sinigaglia et al., 2021) (left of the dash line) and selected markers for epidermis and accessory cells of sensory organs (right of dash line). For each marker we indicate the level of expression per cluster (heat map) and the fraction of cells in each cluster expressing the marker (circle size). Genes associated with molting are upregulated in cell clusters 18, 22 and 28 in the 6-month dataset. Cell clusters 18 and 22 also express markers of epidermis, whereas cluster 28 express one of the markers of sensory organ accessory cells.

**Figure S12.**
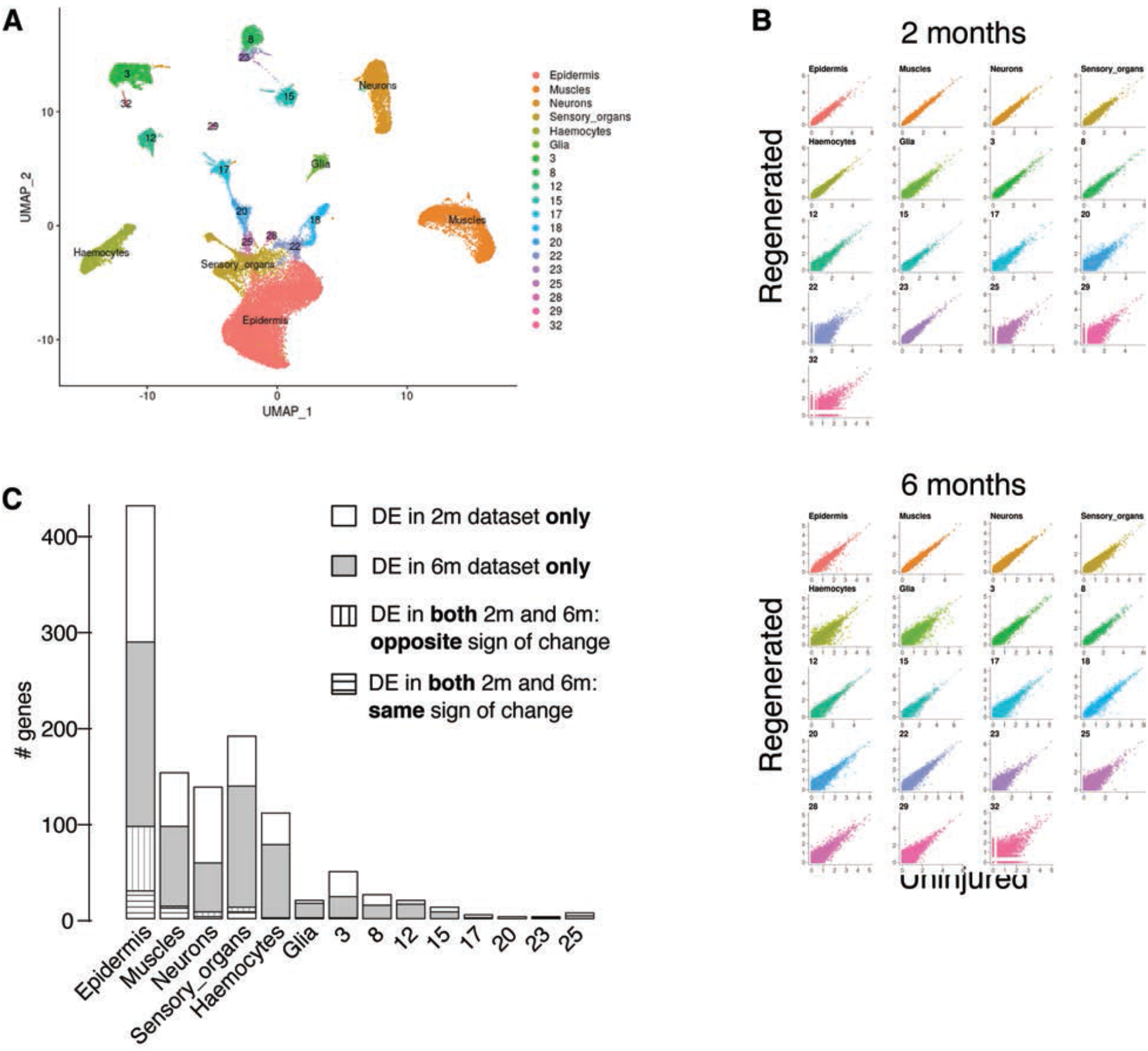
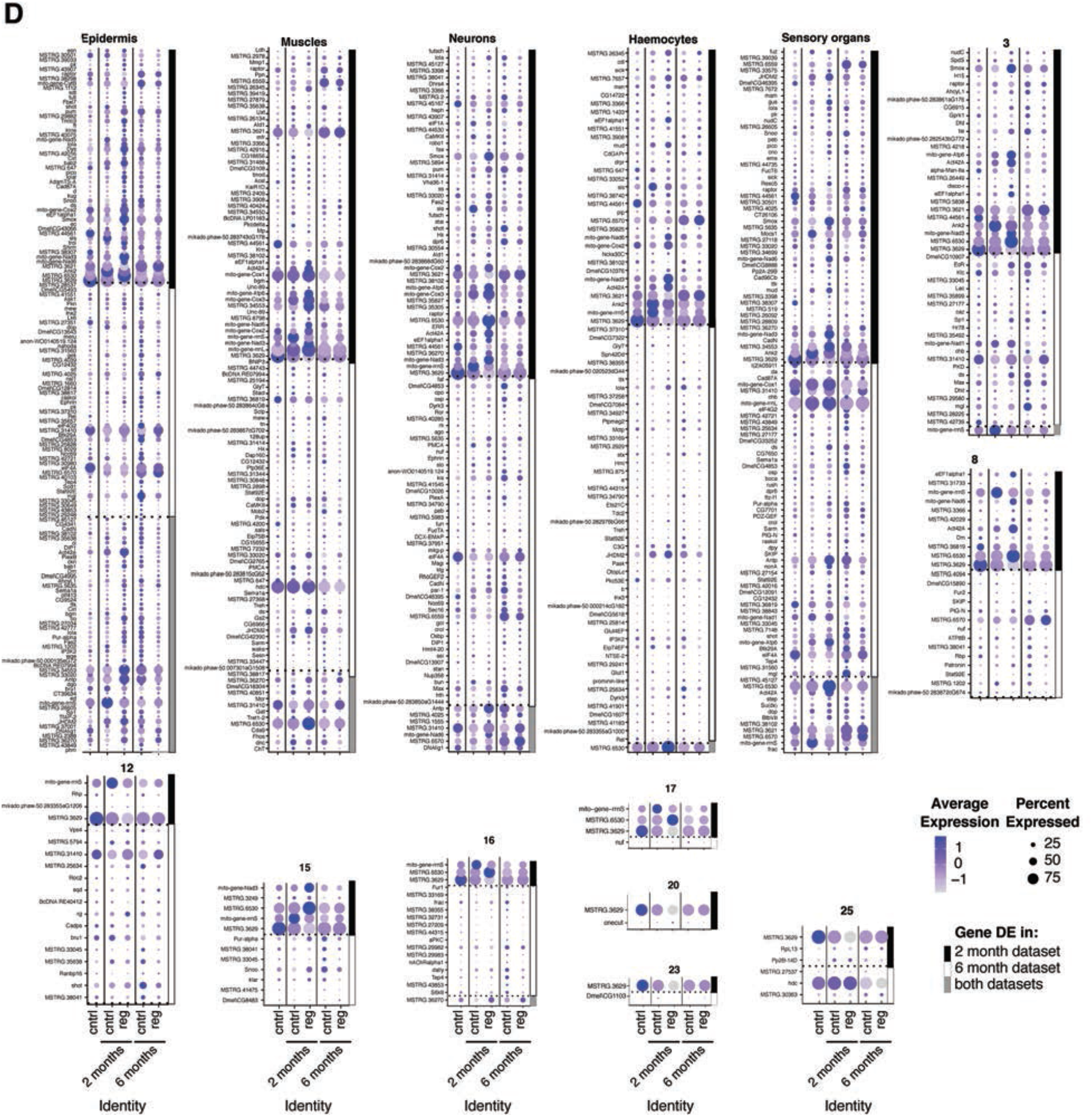
Comparison of transcriptional profiles of regenerated versus uninjured legs. (**A**) UMAP of the five integrated datasets, depicting 40,961 cells clustered based on gene expression (same as in Figure 4A-C, but cell clusters here are shown in distinct colours). (**B**) Comparison of cluster-specific gene expression between control (uninjured) and regenerated samples. For each gene, UMI values were averaged over all the cells assigned to each cluster and were compared between control and regenerated samples separately for the 2-month and 6-month datasets. Cluster-specific gene expression is very similar between regenerated and control samples (Spearman’ s rho of mean ∼0.73 in both datasets). (**C**) Histogram depicting the distribution of genes that are differentially expressed (DE) between control and regenerated samples in the 2-month and 6-month experiments. Most DE genes were found to be differentially expressed in just a single experiment (2-month or 6-month only) or to be differentially expressed in opposite directions in the two experiments. Only a small proportion of DE genes are coherently upregulated or down-regulated in both experiments (horizontal lines), suggesting that a larger share of differential expression is due to experimental variation than to regeneration. (**D**) Dot plot representation of the differentially expressed genes. The top 50 genes that are DE in the 2-month dataset, in the 6-month dataset or in both conditions are displayed (up to 150 genes per cell type). Genes are named with their best reciprocal blast hit in *D. melanogaster* (when available).

**Figure S13.**
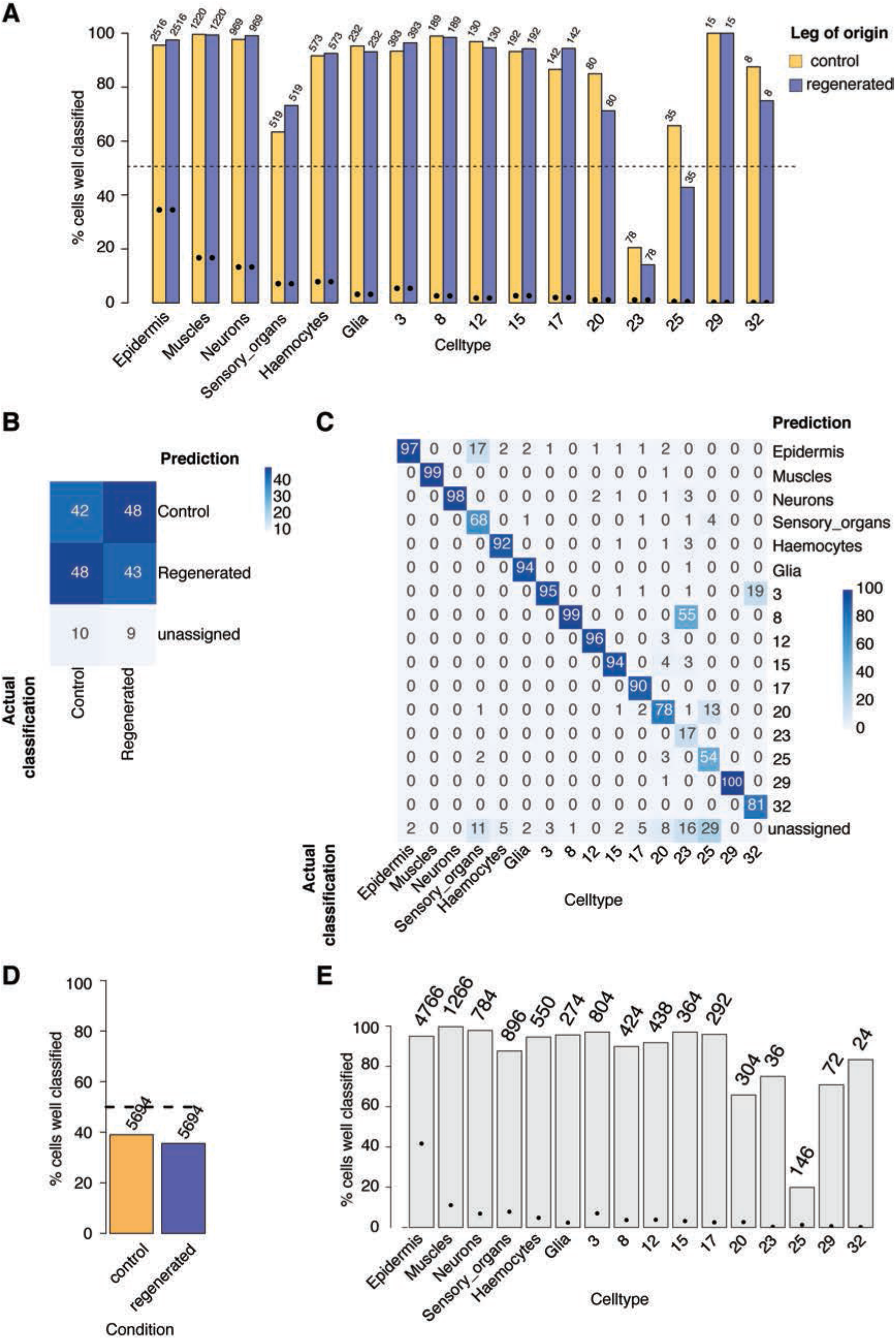
Using machine learning to predict cell type and regeneration status based on the transcriptome. (**A**) Histogram depicting the proportion of cells that are correctly classified with respect to cell cluster/type (detailed version of Figure 4F). (training set = 6-month dataset, testing set = 2-month dataset). (**B**,**C**) Heatmap of the statistics shown in Figure 4F,G. Predicting whether a cell derives from a control or a regenerated leg is no better than a random guess. In contrast, the same machine modelling predicts a cell’ s cluster classification reliably for most cell types. Numbers represent the percentage of cells that are classified in a given category. (**D**,**E**) Same analysis as in Figure 4F,G, but with training and testing sets inverted (training set = 2-month dataset, testing set = 6-month dataset).

**Figure S14.**
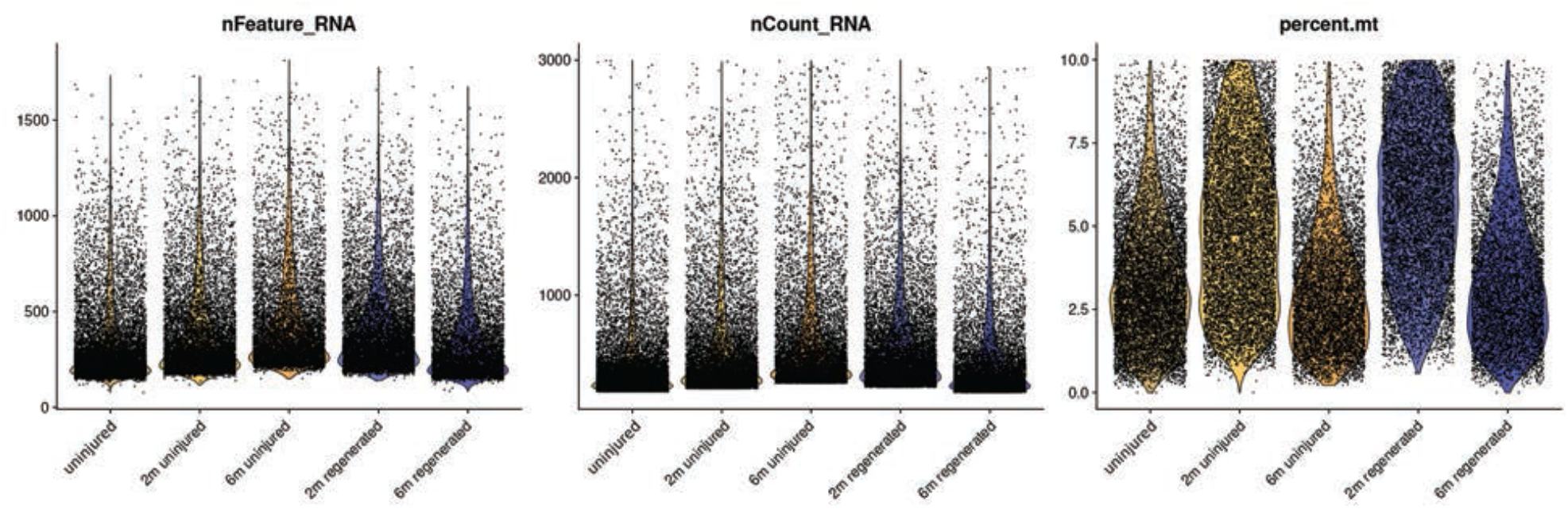
Quality controls for snRNAseq data. Violin plots showing the distribution of the number of genes (nFeatures_RNA), transcripts (nCount_RNA) and percentage of reads mapping to mitochondrial genes (percent.mt) per cell, categorised by dataset.

**Table S1.**
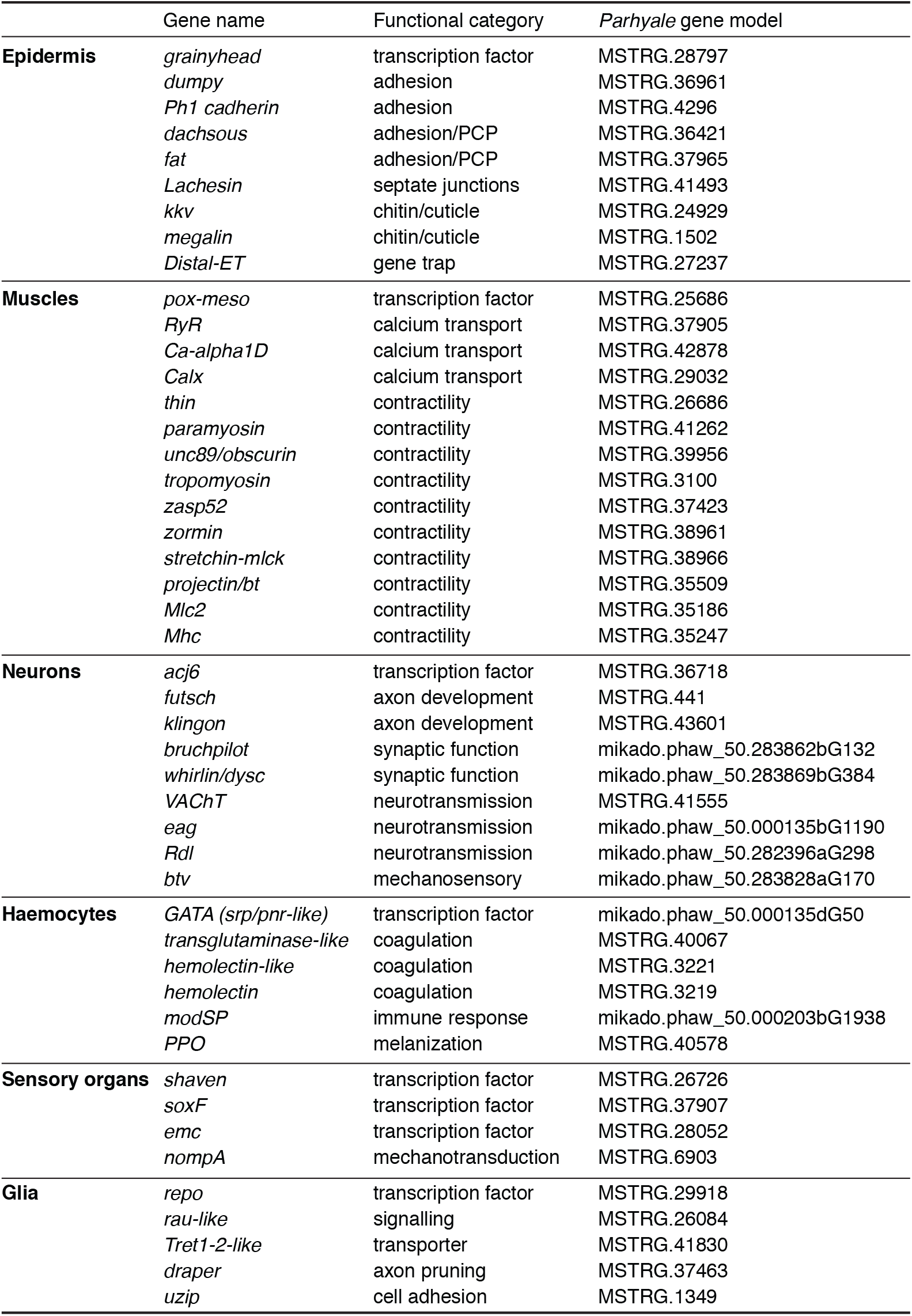
Selected gene markers for the identification of cell types in Figure 3C.

### Supplementary Video

**Caption for Movie S1. Assay of mechanosensory function (.mpeg file)**

Video shows the behavioral response of a *Parhyale* adult to mechanical stimulation of the leg sensory organs. The animal is immobilised on a petri dish and stimulated with a fine metal probe. When the dorsal side of the leg is touched the animal does not react. When the crown and the ventral row setae on the ventral side are touched the animal reacts with a whole-body escape response.

### Supplementary Data

**Caption for Data S1. Setae scored on the distal part of intact and regenerated legs (.csv file)**

List of all the setae scored in T4 and T5 legs and measurements of propodus length. Samples were scored either by scanning electron microscopy (marked SEM in the sample ID) or by laser scanning confocal microscopy. The number of setae in the comb and crown groups could not be resolved with confocal microscopy, therefore these setae were only quantified for the samples imaged by SEM.

**Caption for Data S2. Results of mechanosensory assays on control and regenerated legs (.csv file)**

Records of the responses to mechanical stimulation of the carpus posterior comb, propodus ventral array, and propodus posterior comb (1, response, as shown in Movie S1; 0, no response). Animals were stimulated twice before amputation (before amp), 6 days post-amputation (6 dpa; performed only on the carpus posterior comb), and after the molt following the regeneration (after regen) on both amputated and uninjured control legs.

**Caption for Data S3. Differentially expressed transcripts per cell cluster (.csv file)**

List of transcripts that are differentially expressed in each cluster, including the gene ID, the degree enrichment (fold up- or down-regulation), the statistical significance of enrichment (p-value and adjusted p-value based on Bonferroni correction), and the proportion of cells within the cluster (pct.1) and outside of the cluster (pct.2) containing reads for the transcript.

